# The Phosphate Deprivation Response is Mediated by an Interaction between Brassinosteroid Signaling and Zinc in Tomato

**DOI:** 10.1101/2022.09.21.508943

**Authors:** Gozde S. Demirer, Donald J. Gibson, Xiaoyan Yue, Kelly Pan, Eshel Elishav, Hitaishi Khandal, Guy Horev, Danuše Tarkowská, Alex Cantó-Pastor, Shuyao Kong, Julin Maloof, Sigal Savaldi-Goldstein, Siobhan M. Brady

**Author notes:** Authors have contributed equally (Gozde S. Demirer, Donald J. Gibson).

## Abstract

Phosphate is a necessary macronutrient for basic biological processes, plant growth, and agriculture. Plants modulate their root system architecture and cellular processes to adapt to phosphate deprivation albeit with a growth penalty. Excess application of phosphate fertilizer, on the other hand, leads to eutrophication and has a negative environmental impact. Moreover, phosphate mined from rock reserves is a finite and non-recyclable resource and its levels are nearing complete depletion. Here, we show that *Solanum pennellii*, a wild relative of tomato, is partially insensitive to phosphate deprivation. Furthermore, it mounts a constitutive response under phosphate sufficiency. We demonstrate that activated brassinosteroid signaling through a tomato BZR1 ortholog gives rise to the same constitutive phosphate deficiency response, which is dependent on zinc over-accumulation. Collectively, these results reveal an additional strategy by which plants can adapt to phosphate starvation.

## Introduction

Phosphorus is an essential element that forms the backbone of DNA and RNA, is a key component of cellular energy transport in the form of ATP, part of phospholipids integral to cell membranes, and plays an important role in protein modification and signaling. As such, phosphorus is one of the most common limiting nutrients in crop production and is supplemented as a fertilizer. The global demand for phosphate fertilizer was 43 million tons in 2019 and is expected to increase to 20 million tons by 2030 (López-Arredondo et al. 2014; FAOSTAT 2021). However, phosphate fertilizer production is projected to peak as early as 2033 and known global phosphate rock reserves are likely to be depleted within 50 to 100 years (Cordell and White 2011). Improving crops for increased phosphate acquisition and utilization is a more sustainable solution than supplementing this limiting nutrient. Therefore, it is critical to understand how plants sense phosphate, regulate its uptake and signaling, and acclimate to phosphate limitation.

Plants uptake inorganic phosphate (PO_4_^3-^) through their roots. Due to its low mobility in soil, the majority of inorganic phosphate is present in the topsoil layers (Miguel et al. 2013). One of the adaptive strategies of plants to phosphate deprivation is an adjustment of root system architecture. When starved for phosphate, plants typically develop shallow root systems and increase the formation and elongation of lateral roots to scavenge available phosphate (Williamson et al. 2001; Miura et al. 2011). Additionally, promotion of root hair elongation under phosphate starvation leads to an increased root surface area, improving phosphate uptake (Bates and Lynch 1996). Upregulation of acid phosphatase activity is also a well-known response to P deprivation and is associated with phosphate remobilization (Gao et al. 2017; Hurley et al. 2010; L. Wang et al. 2011, 2014; Zhang, Wang, and Liu 2015).

Mechanisms of phosphate sensing and uptake are well understood in Arabidopsis (Thibaud et al. 2010; Svistoonoff et al. 2007; Ticconi et al. 2009; Song et al. 2016; Bustos et al. 2010; Rubio et al. 2001; Puga et al. 2014; Chien et al. 2018), yet our knowledge in certain crops, such as tomato, is still fragmentary. Hormones additionally mediate plants’ responses to nutrient deficiency. The plant hormone brassinosteroids (BRs) are critical for plant growth and development and various studies established their importance for plant adaptation to environmental stresses (Ackerman-Lavert and Savaldi-Goldstein, 2020; Planas-Riverola et al., 2019; Kono and Yin, 2020; Singh and Savaldi-Goldstein, 2015). However, the role of BR in nutrient foraging only recently emerged (Pandey et al., 2020) and it is functioning to control root length in response to limited phosphate availability in Arabidopsis (Singh et al. 2014, 2018). Activation of the BR signaling cascade begins with BRs binding to the extracellular domain of the BRASSINOSTEROID INSENSITIVE1 (BRI1) and its co-receptor BRI1-ASSOCIATED RECEPTOR KINASE (BAK1). BRI1 and BAK1 activation leads to the inhibition of BRASSINOSTEROID INSENSITIVE2 (BIN2), a Glycogen Shaggy Kinase 3 (GSK3) kinase. In the absence of BR signaling, BIN2 phosphorylates and inhibits various proteins, among them the homologous transcription factors BRASSINAZOLE RESISTANT1 (BZR1) and BRI1-ETHYL METHANESULFONATE-SUPPRESSOR1 (BES1)/BZR2, which are key positive regulators of BR signaling. Phosphorylated BES1/BZR1 translocate from the nucleus to the cytoplasm, while their de-phosphorylation enables binding to promoters of many genes, to transcriptionally coordinate downstream BR responses and other signaling pathways (Wang et al., 2001; Kim and Wang, 2010; Belkhadir and Jaillais, 2015; Nolan et al., 2020). Both transcription factors were isolated as dominant mutant alleles, *bes1-D* and *bzr1-D*, that lack a critical phosphorylation site and hence are constitutively localized to the nucleus where they activate and repress BR genes as well as genes shared with other pathways (Yin et al., 2002; Wang et al., 2002; Oh et al., 2012; Yu et al., 2011).

In Arabidopsis, low phosphate availability reduces the accumulation of BES1/BZR1 in the nucleus, thereby causing changes to root system architecture, including inhibition of the primary root growth and increased lateral root density (Singh et al. 2014). In agreement, *bes1-D/bzr1-1D* have a reduced response to phosphate deprivation. Other physiological responses to phosphate deprivation, an increase in acid phosphatase activity and production of anthocyanins, are also reduced in Arabidopsis *bes1-D*/*bzr1-D* (Singh et al. 2014). Taken together, the constitutively active *bzr1-d*/*bes1-d* transcription factors provide potential targets to block phosphate deprivation responses in other plant species.

Natural variation in crop germplasm often serves as a resource for breeding abiotic and biotic stress-tolerant plants and elucidating their underlying molecular mechanisms. In the case of tomato, *Solanum pennellii* is an inter-crossable wild relative of the domesticated tomato *Solanum lycopersicum*. As *S. pennellii* is adapted to grow in dry, saline, and rocky deserts that are nutrient poor (Nesbitt and Tanksley 2002; Peralta, Knapp, and Spooner 2005; Moyle 2008), its genome has been used to identify loci associated with drought, salt, and pathogen tolerance (Eshed et al. 1992; Gong et al. 2010; Gur et al. 2011; Dehan and Tal 1978; Koca, Ozdemir, and Turkan 2006; Easlon and Richards 2009). Here, we sought to utilize the variation between *S. pennellii* and *S. lycopersicum* to elucidate the molecular mechanisms underlying tomato phosphate deprivation response. *S. pennellii* constitutively mounts physiological and molecular aspects of this response under phosphate sufficiency via a mechanism that incorporates zinc over-accumulation and BR signaling. These data provide a novel mode by which plants can grow under phosphate deficiency and extend our understanding of plant responses to nutrient deficiency.

## Results

### S. pennellii is largely phosphate deprivation insensitive

Given the resistant nature of *S. pennellii* to a variety of abiotic stresses, we tested the hypothesis that *S. pennellii* (LA0716, hereafter referred to as Penn) is phosphate (P) deprivation insensitive relative to the domesticated *S. lycopersicum* cv. M82 (hereafter referred to as M82). We compared the root system architecture (RSA) of these genotypes in sufficient (1 mM KH_2_PO_4_) or limiting P (10 μM trace PO_4_). M82 responded to P deprivation with a decrease in primary root length (p-value < 0.0001) and an overall increase in total lateral root length as measured by a two-way ANOVA (p-value = 0.0007). In contrast, Penn had no significant differences in primary or total lateral root length (Figure 1A and B). In limiting P conditions, elongation of root hairs was observed in M82 (two-way ANOVA p-value < 0.0001) (Figure 1C and D). In contrast, Penn has significantly longer root hairs compared to M82 under sufficient P (p-value < 0.0001), and Penn root hair length reduces in P-deficient conditions (two-way ANOVA p-value < 0.0001) but is still significantly longer than M82 root hairs (two-way ANOVA p-value < 0.0001) (Figure 1C-D).

**Figure 1.**
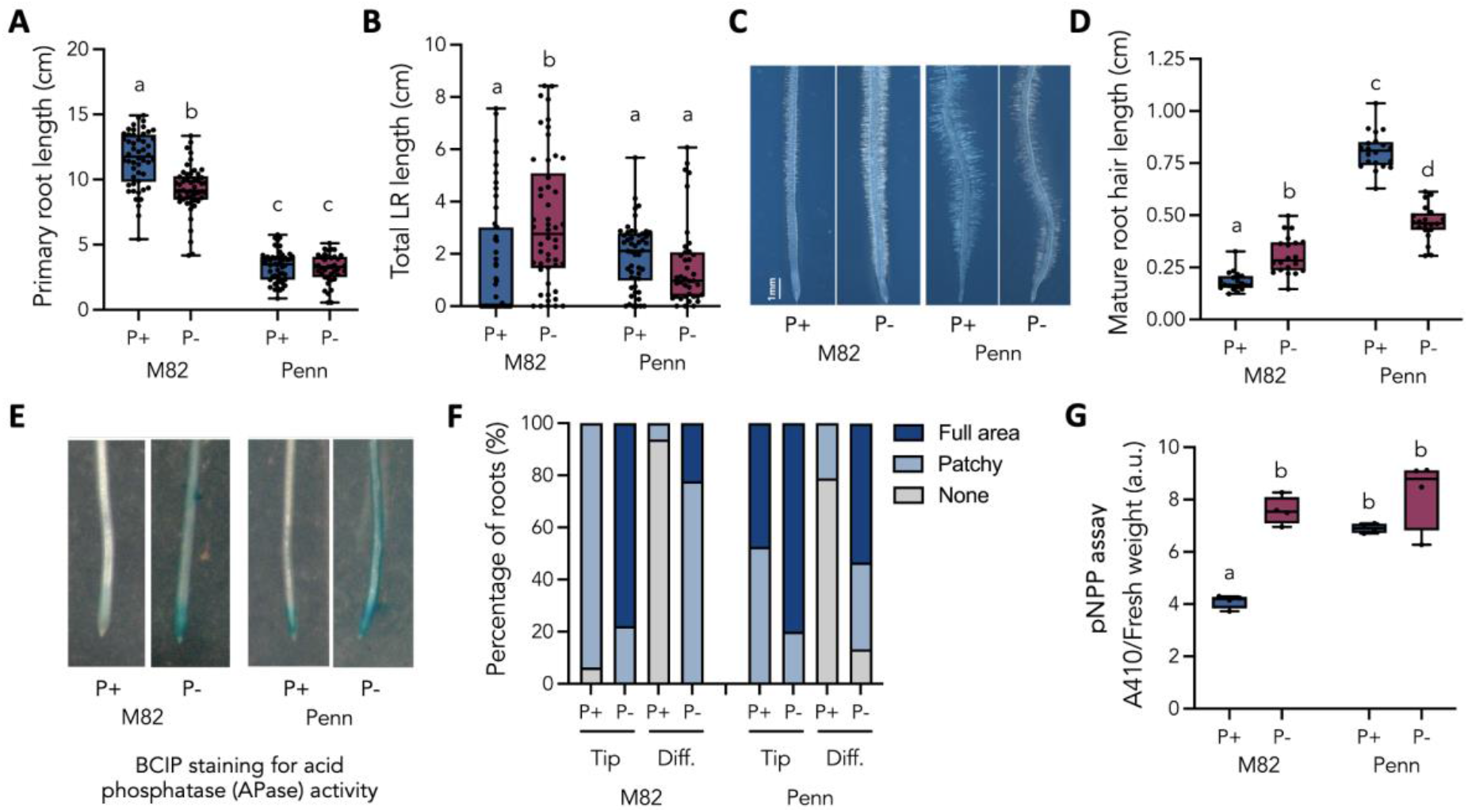
RSA and APase activity response of M82 and Penn to P deprivation. **A)** Primary root length of M82 and Penn (N=50) **B)** Total lateral root length of M82 and Penn (N=50) **C)** Representative root hair images of M82 and Penn **D)** Quantification of mature root hair length for M82 and Penn (N=20) **E)** Representative BCIP APase staining images of M82 and Penn **F)** Qualitative measurement of APase activity in M82 and Penn root tip and differentiation zone **G)** Quantitative p-Nitrophenyl Phosphate (pNPP) cleavage assay of whole M82 and Penn roots (N=4) in P-sufficient and limiting conditions. Letters represent statistically significant differences as determined by a two-way ANOVA and a post-hoc Tukey’s test, p<0.001. Refer to Supp. Fig. 1 for statistical analysis details.

In M82, the increase in acid phosphatase (APase) activity can be visualized using 5-bromo-4-chloro-3-indolyl-P (BCIP) staining in the root tip (meristem and elongation zone) and in the differentiation zone (Figure 1E-F). Qualitative measurement of this activity (complete vs. patchy dye accumulation) indicates that a small amount of APase activity is observed in the M82 root tip under P-sufficient conditions, while in P-deficient conditions, this activity increases to nearly complete staining (Figure 1F). In the M82 differentiation zone, staining is absent in P-sufficient conditions, and present as patchy staining in P-deficiency. On the other hand, Penn has increased APase activity in P-sufficient conditions in the root tip relative to M82 and slight patchy accumulation in the differentiation zone. In response to P-deficiency, this activity becomes nearly complete in the root tip and increases in the differentiation zone (Figure 1E-F). Therefore, although Penn APase activity responds to P-deficiency, it already has increased activity in P-sufficient conditions relative to M82, similar to the root hair elongation in Penn. This trend was corroborated by whole root measurements of p-Nitrophenyl Phosphate (pNPP) cleavage, which is a quantitative colorimetric assay for APase activity (Figure 1G). Here, M82 increases its APase activity in P-deficient conditions (p<0.0001), while Penn has significantly higher activity in P-sufficient conditions compared to M82 (p=0.0011) with a significant genotype by treatment interaction as measured by a two-way ANOVA (Supp. Fig. 1, ANOVA tables associated with figures are in the Supplementary Information).

### S. pennellii morphological insensitivity is reflected in the transcriptome

Plants respond to P deprivation by activating a transcriptional regulatory cascade (Misson et al. 2005; Thibaud et al. 2010; Bustos et al. 2010; Rubio et al. 2001). The reduced RSA and APase activity response in Penn compared to M82 in P deficiency is expected to be accompanied by a similarly dampened transcriptional response. To capture transcript abundance in these conditions and genotypes, the whole root transcriptome was characterized in P-deficient and sufficient conditions in 35S:Translating Ribosome Affinity Purification (TRAP) lines containing FLAG-tagged ribosomes driven by the 35S promoter (Maher et al. 2018). These lines have previously shown to demonstrate excellent correlation with the whole root transcriptome (Maher et al. 2018; Reynoso et al. 2019). Two regions of the root were collected - a one cm segment of the root tip and a two cm section from the middle of the root. This latter tissue contains cells undergoing active lateral root initiation and early stages of lateral root elongation. Differentially expressed genes (DEGs) in each region in response to changes in P concentration were identified using EdgeR (Robinson et al., 2010) with an FDR adjusted p-value of <0.05 (list of DEGs can be found in Supp. Table 1). While M82 has hundreds of genes differentially expressed in both domains of the root (Figure 2A), very few genes are differentially expressed in Penn with most occurring in the middle section of the root in P deficiency as compared to sufficiency (Figure 2B). Minimal overlap of these genes was observed between developmental zones in M82, and no overlap was found in Penn (Figure 2). These changes in gene expression support the dampened response of Penn in response to P deficiency relative to M82.

**Figure 2.**
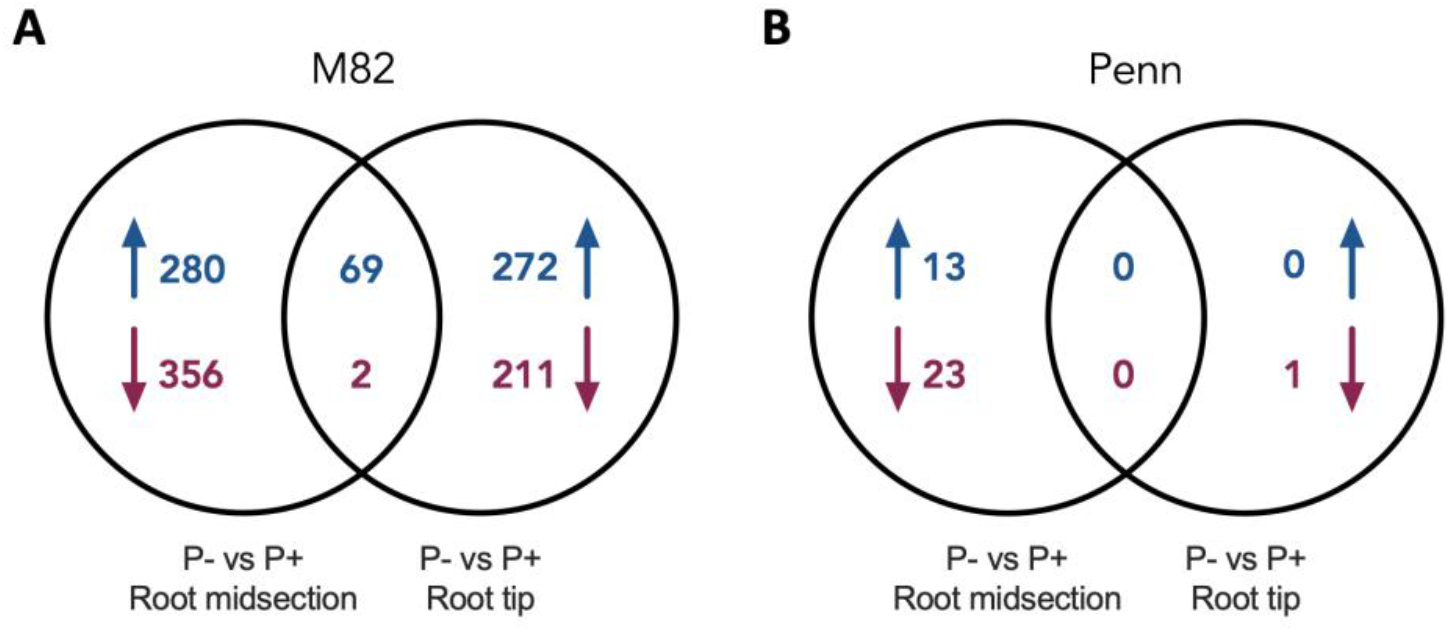
M82 and *S. pennellii* transcriptome response to P-deprivation. **A)** Up- and down-regulated genes in M82 root tip and midsection in P-sufficient and limiting conditions. **B)** Up- and down-regulated genes in Penn root tip and midsection in P-sufficient and limiting conditions.

### Over-accumulation of Zn contributes to ectopic activation of the P deprivation response in S. pennellii

Observed changes in response to P deprivation between these species could be due to underlying differences in P utilization, accumulation, or signaling. For instance, increased P in Penn under P-deficient conditions could explain its insensitivity relative to M82. Signaling could involve interaction between P availability and additional elements, like iron (Fe) and zinc (Zn), the accumulation of which is affected by P availability and vice versa, impacting plant growth rate (Lay-Pruitt et al., 2022).

To test the involvement of ion accumulation levels in the tomato P deprivation response, changes in mineral ion accumulation including P, Fe, and Zn, in the roots and shoots of both species were captured by Inductively Coupled Plasma Mass Spectrometry (ICP-MS). As is typical of plants in P-deficient conditions, both species had lower concentrations of P with no difference between species in either the root or the shoot (Figure 3A and B). A small increase in P in Penn shoot was observed under P-sufficient conditions relative to M82 (two-way ANOVA p-value = 0.0005) (Figure 3B). In low P, Fe is required for inhibition of the primary root growth in Arabidopsis (Svistoonoff et al. 2007). A reduced concentration of Fe in Penn could explain the lack of root growth inhibition in P deficiency. However, the effect of P deficiency on relative Fe accumulation was the same in both species’ root and shoot tissues (Figure 3C and D, Supp. Fig. 3). In contrast, levels of Zn in Penn were nearly four and two times higher in the shoot and root, respectively, compared to M82 in P-sufficient conditions (two-way ANOVA p-value < 0.0001) (Figure 3E and F). Similarly, but to a lesser extent, levels of cadmium (Cd), cobalt (Co), and manganese (Mn) were increased in Penn under P sufficiency (Supp. Fig. 2).

**Figure 3.**
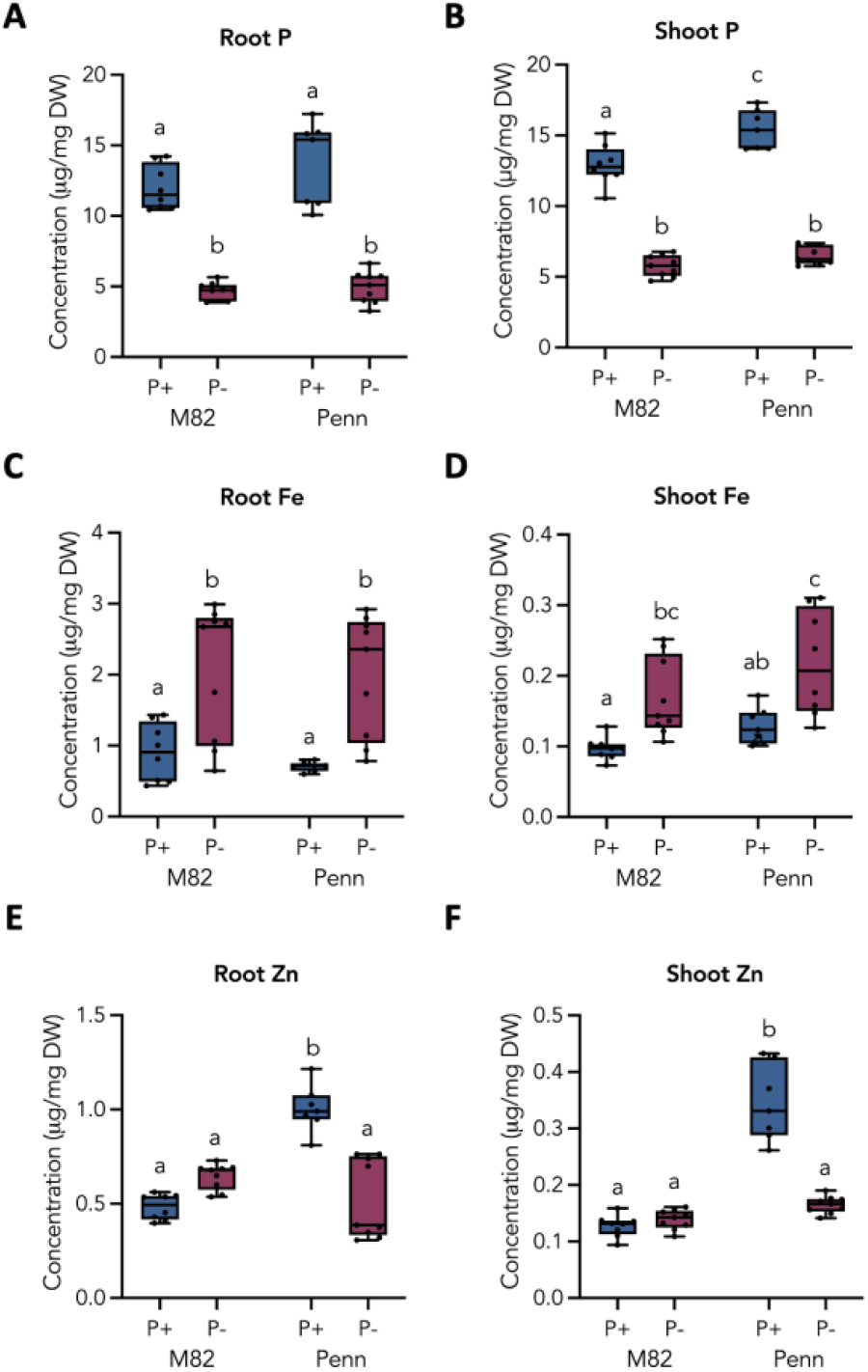
Inductively Coupled Plasma Mass Spectrometry (ICP-MS) of M82 and *S. pennellii* roots and shoots in P-sufficient and limiting conditions. **A)** Root P **B)** Shoot P **C)** Root Fe **D)** Shoot Fe **E)** Root Zn and **F)** Shoot Zn profile of M82 and Penn in P-sufficient and limiting conditions. N = 8, 7, 9, 9 for M82 P+, M82 P-, Penn P+, Penn P-, respectively. Letters represent statistically significant differences as determined by a two-way ANOVA and a post-hoc Tukey test, p<0.03. Refer to Supp. Fig. 3 for F-values.

The ICP-MS results of P and Zn accumulation levels (Figure 3), together with the phenotypic response in P sufficiency and deficiency (Figure 1), suggest that Penn may have constitutive activation of the P deprivation signaling response in P-sufficient conditions. Fe accumulation patterns, which are identical to M82, demonstrate that interaction with Fe is not responsible for these ectopic P-deprivation responses in Penn. Therefore, the increased accumulation of Zn in P-sufficient conditions led to the hypothesis that the high accumulation of Zn may confer the ability of Penn to constitutively activate the P signaling response in the presence of sufficient phosphate.

### Removal of zinc partially rescues the P deprivation insensitivity of S. pennellii

We tested this hypothesis by removing Zn from the plant growth media and observing RSA, root hair, and APase traits in M82 and Penn under both P sufficiency and deficiency. If the excess Zn in P-sufficient conditions is responsible for the ectopic activation of the P-deprivation response, then removal of Zn should suppress this ectopic activation, and result in a differential response in Penn, similar to what is observed in M82. Removal of Zn did not change any traits in M82 compared to Zn-sufficient conditions in either P sufficiency or deficiency (Figure 4A and B). However, in Penn, we observed an increase in total LR length under P deficiency when Zn is removed - a change towards the M82 P deprivation response (two-way ANOVA p-value < 0.0001) (Figure 4C). Similarly, removal of Zn suppressed the increased mature root hair length (two-way ANOVA p-value = 0.0492) (Figure 4D) and APase activity in the root tip (Figure 4E) observed in P-sufficient conditions when Zn was present. In sum, removal of Zn restores Penn traits to those observed in M82 and eliminates activation of the deprivation response in P-sufficiency.

**Figure 4.**
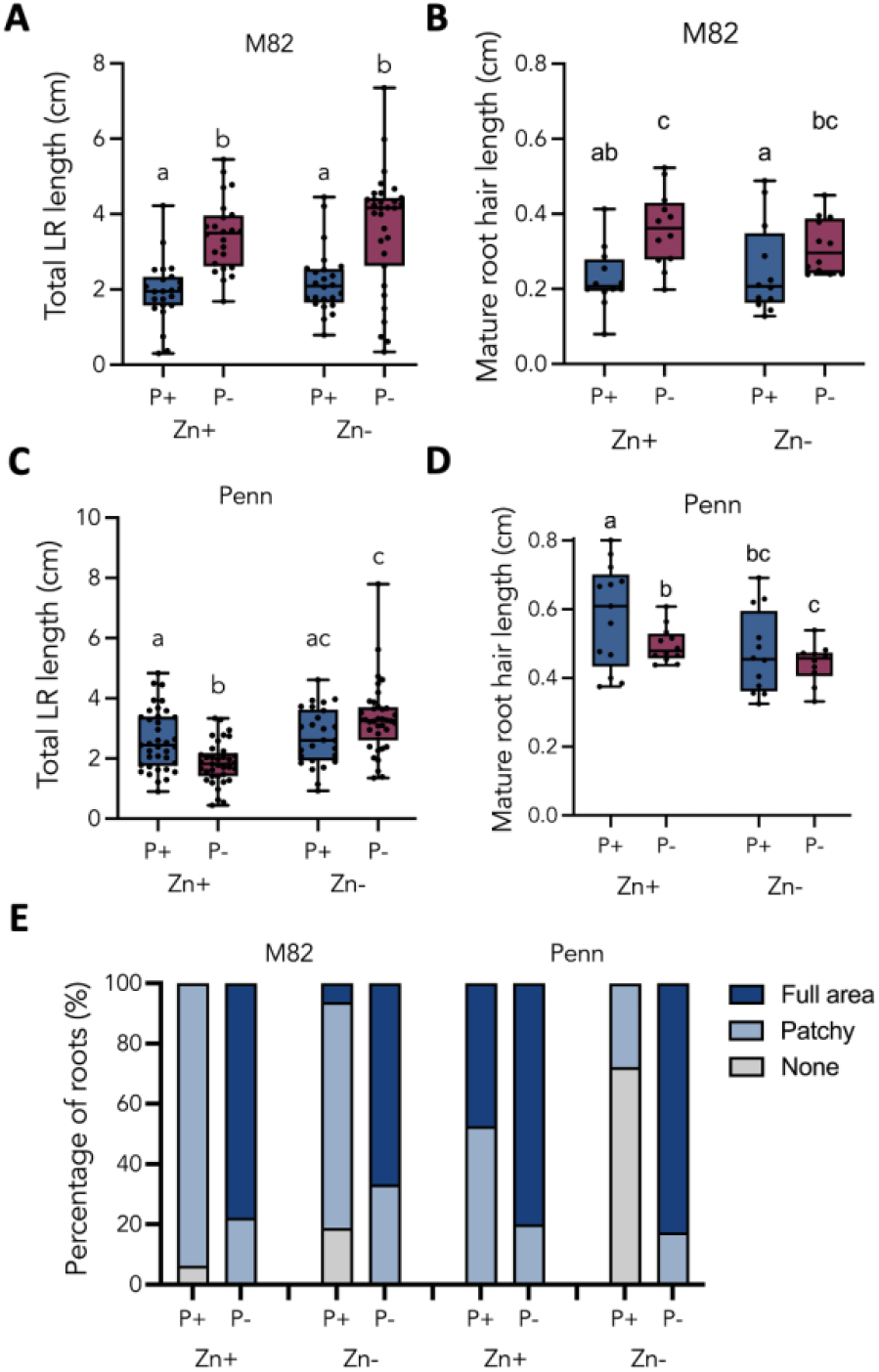
Removal of Zn from growth media. **A)** Total lateral root (N=24) and **B)** mature root hair length responses (N=12) of M82 in P sufficiency and deficiency when Zn is present or absent. **C)** Total lateral root (N=37) and **D)** mature root hair length responses of Penn (N=12) in P sufficiency and deficiency when Zn is present or absent. **E)** APase activity of M82 and Penn in P sufficiency and deficiency when Zn is present or absent (N=21). Letters represent statistically significant differences as determined by a two-way ANOVA and a post-hoc Tukey test, p<0.01. Refer to Supp. Fig. 4 for F-values.

### M82 and Penn differentially accumulate and respond to brassinosteroid levels

Given the interaction between BR and Fe in mediation of the Arabidopsis P deprivation response (Singh et al. 2014, 2018), we hypothesized that in tomato BR levels interact with P and Zn. We first investigated whether there is a difference in BR sensitivity and amounts in M82 and Penn. Penn has increased sensitivity to brassinolide (BL, the most active BR in plants) as shown by a significant reduction of root length relative to no change in M82 upon BL treatment (Figure 5A). In contrast, when the BR synthesis inhibitor, brassinazole (BRZ) (Asami et al., 2000), was added, the Penn root length stayed the same whereas M82 root length decreased substantially (Figure 5B). These results demonstrate that Penn has increased sensitivity to BR application and accordingly reduced sensitivity to BRZ compared to M82, suggesting that Penn may have higher endogenous levels of BR.

**Figure 5.**
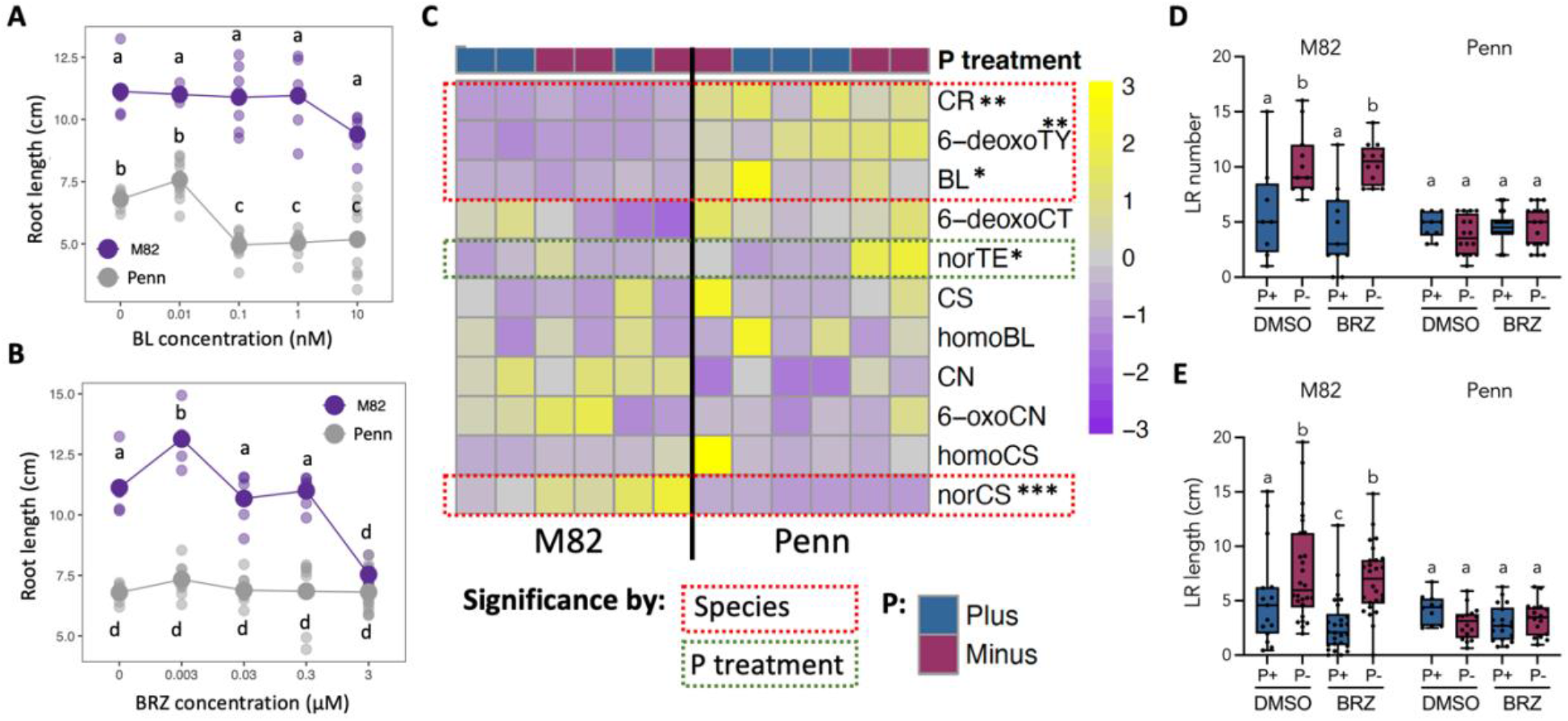
BR sensitivity, amounts, and P interactions in M82 and Penn. **A)** M82 and Penn primary root length measurement upon BL and **B)** BRZ treatment. **C)** BR levels in roots of M82 and Penn in P-sufficiency and deficiency. * denotes statistical difference, red box refers to significance by species, whereas green box refers to significance by P treatment. Shown are: Campesterol (CR), 6-deoxotyphasterol (6-deoxoTY), brassinolide (BL), 6-deoxocathasterone (6-deoxoCT), 28-norteasterone (norTE), castasterone (CS), 28-homobrassinolide (homoBL), campestanol (CN), 6-oxocampestanol (6-oxoCN), homocastasterone (homoCS) and 28-norcastasterone (norCS) **D)** LR number and **E)** LR length of mock and BRZ-treated (3 µM) M82 and Penn plants. Letters represent statistically significant differences as determined by a two-way ANOVA and a post-hoc Tukey test, p<0.05. Refer to Supp. Fig. 5 and 6 for statistical analysis details.

We next tested the influence of BRs in two ways: first, whether the abundance of BRs and their response to P levels are similar or different in roots of M82 and Penn, and second, whether the P-deprivation insensitive response of Penn is relieved upon BRZ application. The levels of BRs were measured in P-sufficient and -deficient conditions in M82 and Penn (Figure 5C). Of those with detectable levels, only nor-TE changed abundance in response to P in both species (adj. p-value <0.05), and four BR types showed species-dependent changes in abundance (CR, 6-deoxyTY, BL, norCS), with three of them being higher in Penn compared to M82 (CR, 6-deoxyTY, BL, adj. p-value <0.05). Higher levels of BL in Penn roots as compared to M82 agrees with Penn’s hypersensitivity to the hormone (Figure 5B). Our next hypothesis to be tested was that this underlying difference in BRs play a role in the phosphate deprivation insensitive response (no change in lateral root number and lateral root length) observed in Penn. Addition of 3 µM BRZ to both M82 and Penn resulted in a slight sensitization increase in M82 with respect to LR length (mixed ANOVA p-value = 0.0176), but no change in the P-deprivation insensitivity observed in Penn (Figure 5D and E). Thus, another factor must be at play in this phosphate-deprivation response insensitivity of Penn.

### BR signaling regulates the phosphate deprivation response in tomato

As BR levels are not responsible for the insensitivity to P deprivation in Penn, we tested if BR signaling, via activated BES1/BZR1, contributes to this lack of response. To test whether constitutive BZR1/BES1 activity blocks the root response to low P in tomato, as in Arabidopsis (Singh et al 2014, 2018), we identified BES1/BZR1 homologs in tomato. *Solyc12g089040*.*1 and Solyc04g079980*.*2* are in a well-supported clade including *AtBZR1 (At1g75080)* (Supp. Fig. 7A). Their functional homology was tested by synthesizing mutations in these two tomato genes to recapitulate the dominant mutation in the *Atbzr1-D* allele. Point mutations were designed to convert a proline to a lysine (Solyc04g079980 - P243->L; Solyc12g089040 - P251->L), and thus remove a likely phosphorylation site. These mutant alleles, fused to the *35S* promoter, were transformed into tomato (*S. lycopersicum* cv. M82) using *Rhizobium rhizogenes* transformation (Ron et al. 2014). Two independent lines with increased expression of each putative *BZR1* ortholog were selected for further analyses (Supp. Fig. 7B). Expression of BZR1 downstream targets, such as *BR6OX2, DWF4, CYP90D1*, and *CPD*, are decreased in *Atbzr1-D* relative to wild type (He et al. 2005). The most likely tomato orthologs of these downstream genes (Zerbino et al. 2018) were identified (*DWF4, Solyc02g085360; CYP90D1, Solyc03g121510; BR6ox2, Solyc02g065750; CPD, Solyc06g051750*) (Wang et al., 2019), and their expression was monitored by real time quantitative PCR in the mutant lines relative to wild type *R. rhizogenes* transformed tomato roots. All four genes had significantly decreased expression in a *Solyc12g089040D*-expressing root, and three of these genes (*CPD, CYP90D1* and *DWF4*) had decreased expression in a *Solyc0460799800D* expressing root (Supp. Fig. 7C). *Solyc04g079980* was named *Slbzr1a-D* and *Solyc12g089040* was named *Slbzr1b-D*, which supports previous observations in tomato (Jia et al. 2021; Yanling Yin et al. 2018; Wang et al., 2019). These same dominant allelic constructs were transformed into M82 using *Agrobacterium tumefaciens* and stable lines were generated.

The dominant mutation found in *Atbzr1-D* renders the plant insensitive to BRZ (Wang et al. 2002; Yin et al., 2002) Two independent lines each from transgenic plants ubiquitously expressing either *Slbzr1a-D* or *Slbzr1b-D* were treated with 5 µM BRZ (BRZ+) or DMSO control (BRZ-), and hypocotyl length of dark-grown plants relative to the M82 was measured. Hypocotyl length of M82 is highly sensitive to BRZ (Figure 6A and B). In comparison, both lines of *Slbzr1a-D* and *Slbzr1b-D* mutants were significantly less sensitive to BRZ (Figure 6A-B for *Slbzr1a-D* line a10, and Supp. Fig. 9 for *Slbzr1a-D* line a7, and *Slbzr1b-D* lines b6 and b22).

**Figure 6.**
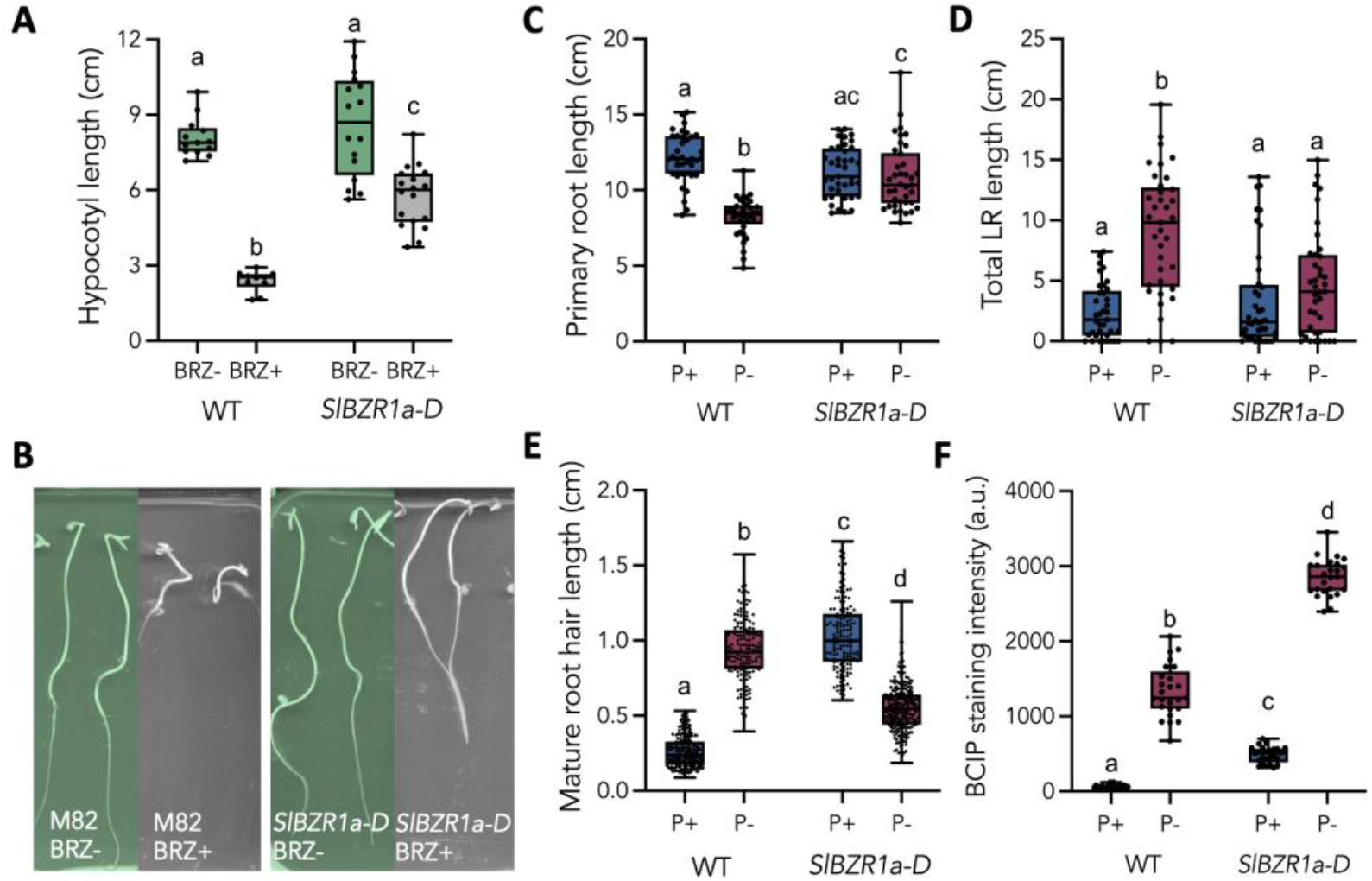
Characterization of the *Slbzr1a-D* mutant (line a10). **A)** Hypocotyl length of dark-grown M82 WT and *Slbzr1a-D* in the absence (DMSO) and presence of 5 µM BRZ (N=10). **B)** Representative photos of dark-grown M82 WT and *Slbzr1a-D* in the absence (DMSO) and presence of 5 µM BRZ. **C)** Primary root length of M82 WT and *Slbzr1a-D* (N=40) **D)** Total lateral root length of M82 WT and *Slbzr1a-D* (N=40) **E)** Quantification of mature root hair length of M82 WT and *Slbzr1a-D* (N=200) **F)** Whole root APase activity via BCIP staining quantification of M82 WT and *Slbzr1a-D* in P-sufficient and limiting conditions (N=25). Letters represent statistically significant differences as determined by a two-way ANOVA and a post-hoc Tukey test, p<0.02. Refer to Supp. Fig. 8 for F-values.

We next tested the hypothesis that BR signaling via *Slbzr1-D* is involved in the M82 response to P deficiency. Indeed, *Slbzr1a-D* is insensitive to deficient P relative to M82 for primary root and total lateral root length (Figure 6C and D for line a-10 and Supp. Fig. 10 for line a7). A similar trend was observed for root hair elongation in *Slbzr1a-D* (Figure 6E) (Figure 1C). Root hairs were longer in P-sufficient conditions relative to M82, while root hair length decreased in response to P-limiting conditions (Figure 6E). *Slbzr1a-D* showed increased APase activity, as measured by BCIP staining, in both P-sufficient and deficient conditions relative to M82 (Figure 6F). A similar RSA response to P-limiting conditions was observed in *Slbzr1b-D* transgenic plants (Supp. Fig. 11), however, plant to plant variation of response was higher. Therefore, for subsequent studies, *Slbzr1a-D* (line a-10) was used as an *Atbzr1-D* homolog.

The phenotypic data presented thus far demonstrate that BR signaling *via* SlBZR1 recapitulates many of the constitutive phosphate deprivation responses observed in Penn. We compared amino acid sequences of *S. pennellii* and *S. lycopersicum* BZR1a (*Sopen04g033560* vs. *Solyc04g079980*) and BZR1b (*Sopen12g030940* vs. *Solyc12g089040*) to investigate whether a mutation at a phosphorylation site of Penn’s BZR1 could explain its P deprivation insensitivity. Even though there are several amino acid differences between the two tomato proteins, the potential phosphorylation domains are fully conserved between *S. pennellii* and *S. lycopersicum* (Supp. Fig. 12).

### BR signaling interacts with Zn to determine the tomato P response

Removal of Zn was sufficient to relieve many aspects of P deprivation insensitivity and ectopic starvation response in P-sufficient conditions in Penn (Figure 4C and D). Next, we sought to elucidate whether BR levels interact with Zn in determining the P response in M82. For this, we tested whether Zn level increase caused by P deprivation was observed upon BL or BRZ treatment in 3 independent biological experiments. As expected, roots of plant grown under P deprivation had lower P levels. No interaction was observed between P accumulation and BL/BRZ treatment (Figure 7A). However, a significant interaction was observed between P availability and BL/BRZ treatment on Zn levels (mixed model ANOVA p=0.00062). Only in response to P-deprivation, BRZ significantly elevated the accumulation of Zn and BL showed the opposite trend in all experiments, although its effect was not significantly different than the DMSO control (Figure 7B). Given that *Slbzr1a-D* fully recapitulates the Penn response in P-sufficient conditions, we anticipated that Zn levels would also increase in this genotype under P sufficiency. Consistent with the phenotype of *Atbzr1a-D, Slbzr1a-D* had no changes in P accumulation in either the root or the shoot relative to M82 (Figure 7C and D). No differences in Fe within the root or the shoot in P sufficiency were observed. Fe concentrations rose in both the root and the shoot in P deficiency, although the magnitude of this change was less in the root of *Slbzr1a-D* (Figure 7E and F). As predicted, Zn was increased in P sufficiency in *Slbzr1a-D* in both the root and the shoot, with the expected changes in P and Fe as in M82 (Figure 7G and H). We also observed an overaccumulation of Zn in Arabidopsis *atbzr1-D* roots in P-sufficiency (Supp. Fig 13F).

**Figure 7.**
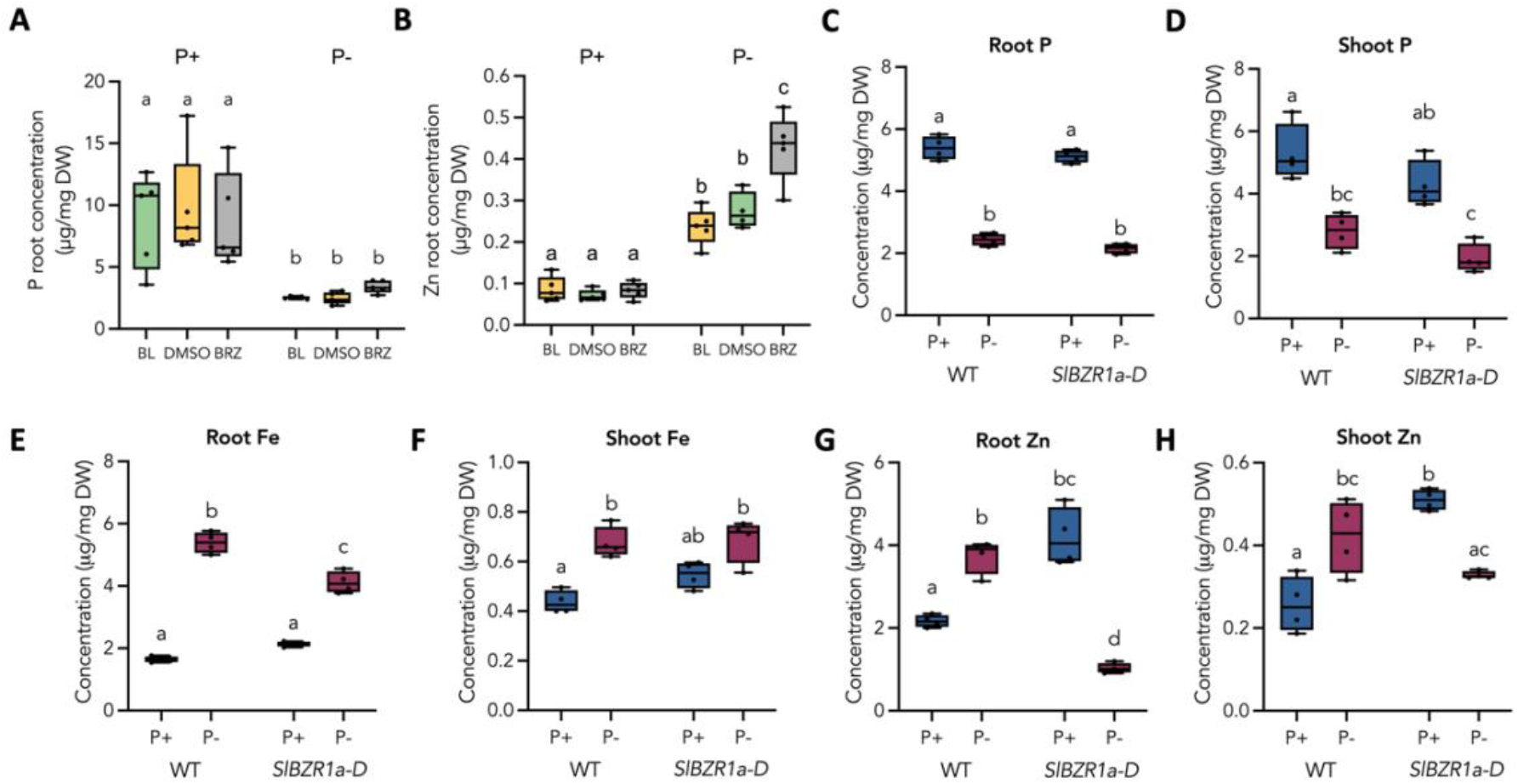
ICP-MS of M82 WT and *Slbzr1a-D* roots and shoots in P-sufficient and P-limiting conditions. **A)** BL/BRZ and P-treatment effect on M82 P and **B)** Zn accumulation measured by ICP-MS. **C)** Root P **D)** Shoot P **E)** Root Fe **F)** Shoot Fe **G)** Root Zn and **H)** Shoot Zn profile of M82 and *Slbzr1a-D* (line a10) in P-sufficient and P-limiting conditions. N= 6 for Panel A and B and N=4 for all other plots. Letters represent statistically significant differences as determined by a two-way ANOVA and a post-hoc Tukey test, p<0.01. Refer to Supp. Fig. 13 for other replicates and statistics and Supp. Fig. 14 for F-values.

Removal of Zn was sufficient to restore APase activity and root hair length in Penn to those observed in M82 (Figure 4). Given the phenocopy of *Slbzr1a-D* to Penn, we next tested if increased Zn accumulation is responsible for some of *Slbzr1a-D*’s distinct P-response perturbations relative to M82. We removed Zn from the media in which plants were grown and observed RSA, mature root hair length, and APase activity of *Slbzr1a-D* and M82 in P sufficiency and deficiency (Figure 8). Primary root length and total lateral root length were now P deprivation responsive in *SlBZR1a-D*, as observed in M82 (Figure 8A,B and E,F). While mature root hair length was not completely restored to M82 levels in *SlBZR1a-D*, they were now responsive to phosphate-deprivation (Figure 8C and G). In addition, increased APase activity levels in *Slbzr1a-D* in P-sufficient conditions were returned to WT levels when Zn was removed from the growth media (Figure 8D and H). These combined results of Penn and *Slbzr1-D* validate our hypotheses that in tomato i) the BR signaling pathway interacts with the P availability response, ii) Penn and *Slbzr1-D* have an ectopic P starvation response, and that iii) Zn controls or at minimum influences the P availability response in tomato.

**Figure 8.**
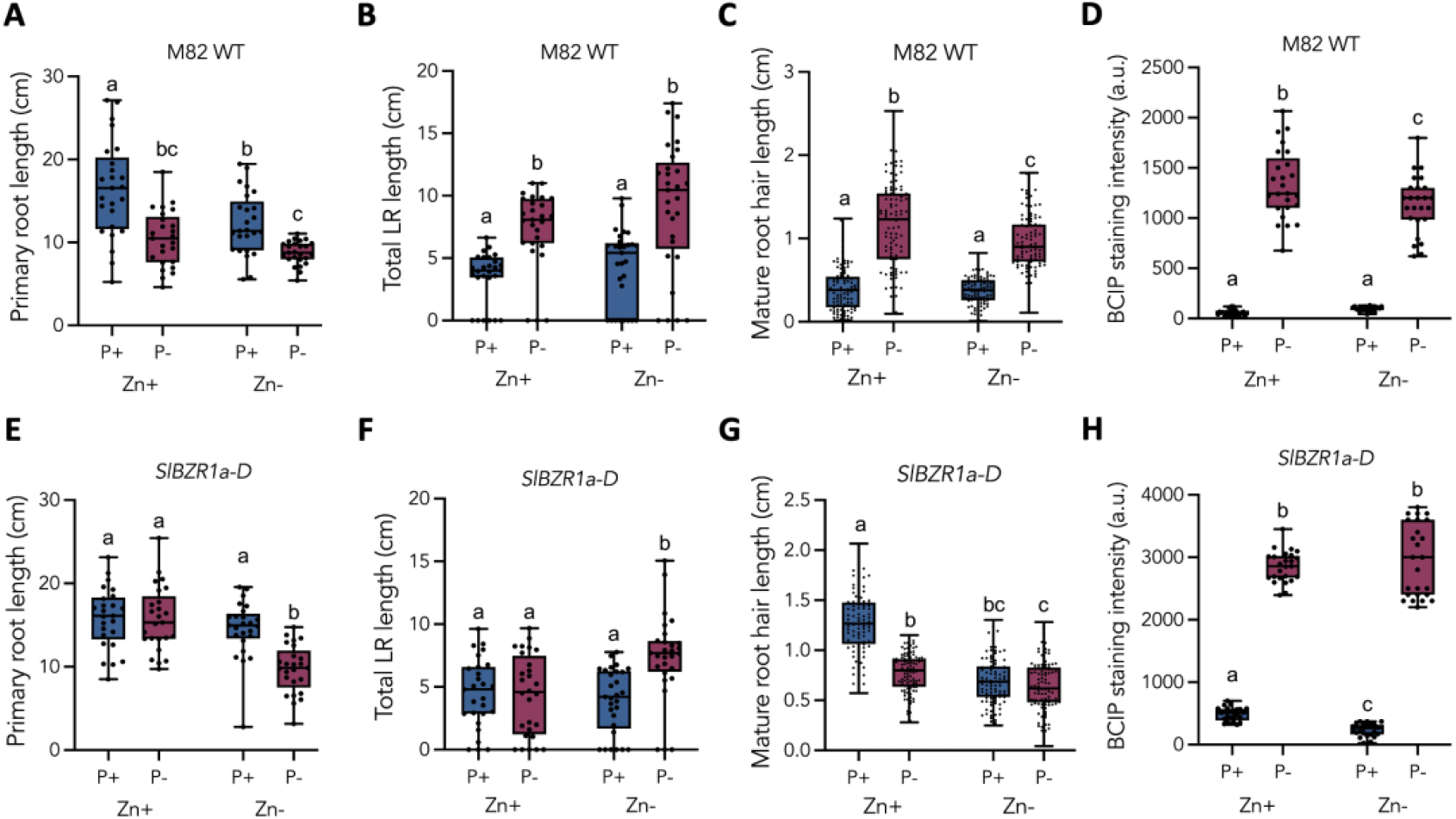
Zn removal from the P-sufficient or P-deficient media of M82 WT and *Slbzr1a-D* plants. **A)** Primary root length **B)** Total lateral root length **C)** Mature root hair length and **D)** APase activity of M82 WT in P-sufficient or P-deficient conditions when Zn is present or absent. **E)** Primary root length **F)** Total lateral root length **G)** Mature root hair length and **H)** APase activity of *Slbzr1a-D* in P-sufficient or P-deficient conditions when Zn is present or absent. Letters represent statistically significant differences as determined by a two-way ANOVA and a post-hoc Tukey test, p<0.01. Refer to Supp. Fig. 14 for F-values.

## Discussion

*S. pennellii* is well known for its resistance to a variety of abiotic stresses (Eshed et al. 1992; Gong et al. 2010; Gur et al. 2011; Dehan and Tal 1978; Koca, Ozdemir, and Turkan 2006; Easlon and Richards 2009), and our results demonstrate that phosphate deprivation is no exception. The phenotypes observed in this wild species are reminiscent of those observed in Arabidopsis *bzr1*-D, where primary root length, lateral root density, and acid phosphatase activity are insensitive to P deficiency relative to wild type (Figure 1) (Singh et al. 2014, 2018). As in Arabidopsis, P deficiency results in a major transcriptional response in M82, while the lack of a morphological response in *S. pennellii* was mirrored in a minimal transcriptional response (Figure 2). Some differences in traits were noted between Arabidopsis and tomato relative to *S. pennellii* that led to the hypothesis that the *S. pennellii* insensitivity is due to an already activated P deprivation response in P-sufficient conditions. In *S. pennellii*, these include decreased root length, increased acid phosphatase activity, and elongated root hairs in P sufficiency (Figure 1).

These observations could additionally be explained by other mechanisms. For instance, the root of *S. pennellii* is shorter than that of M82 and this is a multigenic trait (Ron et al. 2013; Toal et al. 2018), and as shown here, also involves higher BR sensitivity. Arabidopsis root hair elongation is also controlled by the plant hormones, ethylene and auxin (Feng et al. 2017; Pitts, Cernac, and Estelle 1998; Bhosale et al. 2018). Other species-level differences exist between Arabidopsis and M82 in terms of the “classical” phosphate deprivation response. In Arabidopsis, upon P deficiency, the shoot has increased Fe levels, while the root has decreased Fe (Singh et al. 2014). In M82, both the shoot and the root have increased Fe in phosphate deficiency (Figure 3). The BR, nor-CS, is decreased in P deficiency in Arabidopsis, but not in tomato. Nor-TE is the only measured BR that changes levels upon phosphate deficiency in tomato (Figure 5).

Our ultimate hypothesis for the mechanism of *S. pennellii’s* phosphate deficiency insensitivity is an interaction between enhanced BR signaling and increased Zn accumulation. The first experiment that suggested this role of BR was that *S. pennellii* root length is hypersensitive to the addition of exogenous BL and in accordance, hyposensitive to reduction in BR levels (as employed by BRZ, Figure 5A-B). Furthermore, BR precursors (CR, 6-deoxo TY) and the active hormone BL are higher in *S. pennellii* in both P-sufficient and -deficient conditions, in line with its hypersensitivity to the hormone (Figure 5C). Addition of BRZ moderately sensitized the response of *S. lycopersicum* to P deprivation (RSA, Figure 5D-E) as well as the extent of Zn elevation under these conditions (see below, Figure 7A). However, BRZ did not affect *S. pennellii*, indicating a more prominent sensitized signaling. Evidently, the *Slbzr1a-d and Slbzr1b-d* mutants recapitulate the *S. pennellii* observations (Figure 6).

Like the Arabidopsis *bzr1-D* and *bes1-D*, the *Slbzr1b-D* mutant displayed a similar insensitivity to BRZ as *Slbzr1a-D*. As such, *BZR1b* and *BZR1a* are likely *BES1/BZR1* orthologs (Figure 6, Supp. Fig. 11), and we hypothesize that these genes may interact redundantly to control the P deprivation response. *BZR1A* or *BZR1B* could therefore be candidates for the insensitivity observed in *S. pennellii*. However, there are no changes in the phosphorylation domain of BZR1 or the locus where the bzr1-D mutation derives in both *S. pennellii BZR1A* or *BZR1B* (Wang et al. 2002; Yin et al., 2005) (Supp. Fig. 12).

An additional mechanism to explain the P deprivation insensitive phenotype in both *S. pennellii* and *Slbzr1a-D* is that the P deprivation-associated modulation of root system architecture is blocked by elevating endogenous P and decreasing Fe content to compensate for the low P. Using ICP-MS, these predicted changes in P and Fe were not observed, and instead it was evident that Zn levels were increased in P-sufficient conditions in both *S. pennellii* and the *Slbzr1a-D* mutant (Figures 3,7). Removal of Zn in both genotypes restored many aspects of the phosphate deficiency response (Figures 4,8). Therefore, Zn over-accumulation could be a common mechanism by which the activated P deprivation response in *S. pennellii* and *Slbzr1a-D* occurs. These results complement those that previously demonstrated the molecular link between Zn and P accumulation in Arabidopsis (Khan et al. 2014; Bouain et al. 2019). Interestingly, we also observed an overaccumulation of Zn in *atbzr1-D* roots, as dependent on P availability, the implication of which remains to be explored in Arabidopsis (Supp. Fig 13F). Repression of Zn uptake in *S. pennellii* and *Slbzr1a-D* would conclusively determine if Zn overaccumulation is indeed responsible for their respective P deprivation insensitivity and for activation of the P deprivation insensitive response. As a corollary, increasing Zn levels in M82 should be sufficient to result in P deprivation insensitivity and an ectopic response in P sufficiency. Finally, activation of BR signaling in M82 by an additional mechanism (such as a loss-of-function of the BIN2 ortholog(s)), or generation of reduced BR signaling (such as a gain-of-function mutant in the BIN2 ortholog(s)) (Jianming Li and Nam 2002) in *S. pennellii* should result in P deprivation insensitivity and suppression of P deprivation insensitivity, respectively.

How then does BR signaling determine changes in Zn levels? One candidate is the IRT1 transporter. In Arabidopsis, IRT1 transports Fe, Zn, Cd, Mn and Co (Eide et al. 1996; Korshunova et al. 1999; Vert et al. 2002). While no change in Fe uptake was observed in *S. pennellii* or *Slbzr1a-d*, in addition to Zn, an increase in Cd, Mn, and Co were observed in phosphate sufficiency relative to M82 (Supplementary Figure 2). The tomato ortholog of this transporter is therefore an attractive candidate for this trait. IRT1 specificity can be modulated by mutation of certain amino acids. Replacing the aspartic acid residues at positions 100 or 136 with alanine eliminates transport of both Fe and Mn, while replacement of a glutamic acid residue at position 103 eliminates its ability to transport Zn (Rogers, Eide, and Guerinot 2000). Manipulation of IRT1 via these types of mutations in M82, *S. pennellii*, and *Slbzr1a-d* could further connect the interaction between BR signaling and P deprivation insensitivity.

In conclusion, the tomato wild species *S. pennellii* mounts an ectopic phosphate deprivation response in phosphate sufficiency and is insensitive to phosphate deficiency. This trait is phenocopied in a dominant mutation of tomato *SlBZR1A* and suggests that BR signaling plays a role in the *S. pennellii* response. Zinc overaccumulation is observed in both of these genotypes, and removal of zinc is sufficient to suppress the ectopic and insensitive phosphate deprivation response in both genotypes. Collectively, these data provide an alternative mode by which the phosphate deprivation response is regulated and could be used to breed plants that are better adapted to withstand phosphate deficiency. These findings are particularly important given the detrimental nature of phosphate fertilizer overapplication in agricultural settings and the imminent exhaustion of phosphate mineral reserves.

## Materials and Methods

### Plant Growth

*Solanum lycopersicum* var. M82 WT and *Slbzr1-D*, and *Solanum pennellii* var. LA0716 seeds were sterilized using a treatment of 5 minutes of 70% ethanol, followed by 15 minutes of 50% bleach, and then rinsed five times with sterile water. Seeds were sowed directly on sufficient phosphate MS media, 1 mM KH_2_PO_4_, or limiting phosphate (10 µM of trace phosphate) MS media. MS media with sufficient phosphate contained the following nutrients: 20.4 mM NH_4_NO_3_, 18.8 mM KNO_3_, 4 mM CaCl_2_, 1.25 mM MgSO_4_, 1 mM KH_2_PO4, 0.1 mM FeSO_4_, 0. 1 mM EDTA, 0.1 mM H_3_BO_3_, 0.0994 mM MnSO_4_, 0.048 mM ZnSO_4_, 5 µM KI, 1.036 µM Na_2_MoO_4_, 0.096 µM Co_2_Cl_2_, 0.05 µM CuSO_4_, 0.1 g/L *myo*-insotiol, 0.5 mg/L Pyridoxine HCl, and 0.1 mg/L Nicotinic Acid. Media was bound using Difco Granulated Agar (VWR Catalog Number 90000-786) at 7 g/L. In phosphate limiting media, 1 mM KH_2_PO_4_ was replaced with 1 mM KCl. Limiting phosphate media contained approximately 10 µM phosphate due to trace amounts found in the agar from data presented by Difco. For preparing Zn-deficient media, the same recipe is followed but with complete removal of 0.048 mM ZnSO_4_. Seedlings were grown on plates, (245mm x 245mm x 10mm; Corning ™ catalog number 431272) containing 200 to 250 mL solid media. Seedlings were grown in a controlled environment facility with 12 hours of day and 12 hours of night. The light intensity ranged between 75 and 85 µM per meter squared per second and the red to far-red ratio was approximately 1.7. Unless stated otherwise plants were measured at 9 days after germination.

### Root System Architecture Analysis

Seedlings were sowed directly onto P sufficient or P limiting, or Zn sufficient or Zn starved, media containing agar. Germination was tracked daily and plates were scanned using an Epson Perfection V800 Photo Color Scanner into 24-RGB TIF image files at 600 dpi. RSA was quantified at 9 days after germination for each individual seeding to control for age. Images were processed using the ImageJ plugin SmartRoot (Lobet, Pagès, and Draye 2011) and four root traits were quantified: Primary Root Length, Number of Lateral Roots, Length of Lateral Roots, and Total Root Length. Primary Root Length is the length from the root-hypocotyl junction to the tip of the primary root. Number of Lateral Roots is the number of lateral roots per individual seedling. Length of Lateral Roots is the combined length of all the lateral roots for one seedling. Total Root Length is the combined length of primary and lateral roots for one seedling. Measurements were combined and processed using a custom R script.

### BL/BRZ treatments and RSA study

For hormonal sensitivity assays (Figure 5A,B), seedlings were germinated on 0.5 MS and were transferred after 5 days to media supplemented with increasing concentrations of BL and BRZ (as indicated in the corresponding plots) for additional 7 days. For RSA analysis (Figure 5D,E), seedlings were germinated onto P-sufficient or P limiting and were transferred after 7 days to their corresponding media supplemented with DMSO or 3 µM BRZ and analyzed after an additional 7 days. Images from the scanned plates were analyzed using Fiji to quantify root growth parameters as above.

### Root Hair Measurements

Tomato seedlings were grown on phosphate sufficient or limiting, or Zn sufficient or Zn starved, conditions for between 8 to 10 days after germination. Four root hair traits were quantified using a custom R script and ImageJ measurements for the following traits: “Root Tip to First Hair”, “First Root Hair to Maturity”, “Root Hair Length”, and “Root Diameter”. Using ImageJ, a straight line was made from the root tip until the first root hair was observed. This distance was named “Root Tip to First Hair”. Then, a second straight line was made until the root hairs appeared to reach their mature length. This region was named “First Root Hair to Maturity”. From this point on, a line along the root was taken every 1 mm for a total of 10 mm, at the point of each line, a segment of the root width and root hair length were taken. The average of the 10 points was taken for “Root Hair Length”.

### 5-bromo-4-chloro-3’-indolyphosphate p-toluidine (BCIP) Staining for Qualitative Acid Phosphatase Response

To visualize acid phosphatase activity on tomato roots, 5-bromo-4-chloro-3’-indolyphosphate p-toluidine (BCIP) (Sigma-Aldrich catalog number B6149) staining was performed. BCIP (0.02% w/v) was added to 5 g/L agar solution of ddH_2_O at pH of 5.8 and applied on the tomato roots at a temperature of 40°C with an approximate volume of 50 mL. After the agar solidified (within minutes), images were taken at zero hours and two hours after solidification. Images were taken of the root tip of plants grown in limiting P or sufficient P, and/or in Zn sufficient or Zn starved media depending on the experiment. Roots were qualitatively measured based on one of three levels of staining: No Staining, Patchy or Light, and Full Area in a blinded manner by two different individuals. These qualitative measures were based on visual detection of the images with No Staining referring to root tips with no evidence of stain, Patchy or Light are roots with some staining or a light stain, and Full Area refers to staining across the entire area examined. The root sections investigated were the Root Tip, approximately one mm from the root tip and the Differentiation Zone approximately two to five mm from the root tip.

### Nitrophenyl Phosphate (pNPP) Acid Phosphatase Quantification

Whole root systems were excised from MS plates then weighed and washed lightly in water which was blotted with a paper towel. The root system was then placed into buffer containing 50 mM Sodium Acetate, 1 µM dithiothreitol (DTT), and 6 µM p-Nitrophenyl Phosphate (pNPP) (New England Biolabs Catalog Number P0757S) at 25°C at a pH of 5.0. Measurements were taken at the absorbance A410, 1 hour after placing the roots into the buffer. Four biological replicates and three technical replicates of each biological replicate were performed per genotype per condition. Estimation of activity was determined as Absorbance at 410 nm/Fresh Weight.

### Translating Ribosome Affinity Purification-RNAseq

35S:TRAP construct-containing *S. lycopersicum* var. M82 and *S. pennellii* var. LA0716 containing seedlings (Reynoso et al. 2019) were harvested between eight and ten days after germination. Two tissues were extracted for analysis, the root tip and the root midsection. The root tip comprised a 1 cm segment from the tip of the root while the midsection was approximately 2 cm of tissue halfway between the root-shoot junction and the root tip. RNA libraries were prepared using Breath Adapter Directional sequencing (BrAD-seq) a strand-specific 3-prime RNA-seq library prep protocol (Townsley et al. 2015). A total of 4 biological replicates for each genotype and treatment were used with between 30 and 40 plants in each biological replicate. Libraries were pooled, barcoded, and submitted to the UC Davis DNA Technologies Core and were sequenced on Illumina HiSeq2000 SR50. A custom perl script partitioned the reads into individual libraries based on the barcode and trimmed reads. FastQC (https://www.bioinformatics.babraham.ac.uk/projects/fastqc/) was used to check the quality of each sequenced library. Five of the 32 libraries were excluded from analysis due to poor sequence quality. Reads were aligned to the *S. lycopersicum* reference transcriptome ITAG 3.1) (Fernandez-Pozo et al. 2014) with bowtie 1.2.2 (Langmead, 2010) (http://bowtie-bio.sourceforge.net/index.shtml). Reads were aggregated in count tables using a custom script. Differential gene analysis was performed on normalized read counts using the method of trimmed mean of M-values (TMM) with the calcNormFactors function in edgeR (Robinson et al., 2010) and the limma (Ritchie et al., 2015) Bioconductor packages in R. A linear model was used to determine the effects of developmental stage and phosphate by the developmental state on differential gene expression. *S. lycopersicum* and *S. pennellii* were analyzed separately but used the same pipelines and thresholds for analysis. Multiple testing correction was performed using the ashR package in R. P-values are set below 0.05.

### ICP-MS Elemental Analysis

Seedlings aged eight to ten days after germination were harvested and separated into roots and shoots for fresh weight measurement. Seedlings were then dried at 50°C for 24 hours, measured for their dry weight and then ground into a powder using Zirconia Silica beads (2.3mm diameter, catalog number 11079125z). M82 and Penn samples were analyzed by inductively coupled plasma mass spectrometry (ICP-MS) at the UC Davis/Interdisciplinary Center for Plasma Mass Spectrometry (UCD/ICPMS) to measure the relative mass of 30 elements including P, Fe, and Zn. For each genotype, tissue, and condition, two experimental replicates were performed, and each experiment had four biological replicates. M82 WT and *Slbzr1-D* samples were prepared the same for ICMPS, and measured at the UC Berkeley College of Natural Resources Inductively Coupled Plasma Spectroscopy Facility. For analysis of plants subjected to hormonal treatment (Figure 7), seedlings were germinated in pots containing perlite, watered with P sufficient (1 mM) or P limiting (20 µM) MS media. After 7 days, the seedlings were transferred to new pots and were watered with the corresponding P media supplemented with DMSO (mock), BRZ (3µM) or BL (10nM), and grown for additional 14 days. Three independent biological replicates were performed, of 3-5 plants per treatment. Analysis (using HR ICP-OES PQ 9000, Analytik Jena, Germany) was performed at the Faculty of Civil and Environmental Engineering, Technion or at the Interdepartmental Equipment Facility, HUJI.

### *Rhizobium rhizogenes* “Hairy Root” Transformation

Transformation with *Rhizobium rhizogenes* was performed as previously described (Ron et al. 2014). Sterilized tomato cv. M82 seeds were sown in magenta boxes containing 4.3 g L^-1^ Murashige and Skoog (MS) medium (Caisson; catalog no. MSP01-50LT) with 0.5 g L^-1^ MES, 10 g L^-1^ Sucrose, 10 g L^-1^ agar (Difco; catalog no. 214530) at pH 5.8. Seedlings were grown under 16H Day / 8H Night conditions at 22°C to the point at which the cotyledons are fully expanded (usually 7 to 10 days after germination). Competent *R. rhizogenes* were electroporated using the desired construct and plated on plates containing MG / L bacterial growth media (Caisson; catalog no. MQP04-1LT) with 50 mg L^-1^ kanamycin as antibiotic selection. Plates were grown for three days at 28°C, individual colonies were selected, and the correct insert was confirmed via PCR-based genotyping with a gene-specific primer. Selected *R. rhizogenes* colonies were inoculated in 10 mL of liquid Caisson medium (MG/L) at 28°C and shaken at 200 rpm for 24 hours.

Approximately 40 cotyledons per construct were cut and placed into liquid half strength MS. Cotyledons were then incubated in the *R. rhizogenes* for 20 minutes and then transferred to half strength solid MS with 3% sucrose and 7 g L^-1^ agar to incubate for three days, in the dark at 22°C. After three days the inoculated cotyledons were transferred to a selection media containing half strength solid MS with 3% sucrose, 200 mg/L cefotaxime, and 50 mg/L kanamycin. The successfully transformed cotyledons then produced roots over a two-week period. Each root grown from a cotyledon is a unique transformation event and these individual transformation events were subcloned every two to four weeks into a new plate to ensure each testing of multiple biological replicates of each insertion line.

### Generation of Dominant Mutant Alleles in SlBZR1 Putative Orthologs

Pairwise genome alignments between these proteins were generated using LastZ and BLAST which were chained together to result in a lastZ-net alignment (Zerbino et al. 2018). DNA sequences for these protein coding regions were synthesized by Bionexus™ into a GATEWAY™ compatible vector pUC57 with point mutations made to convert a proline to a lysine in Solyc04g079980 (P243-> L) and Solyc12g089040 (P251 -> L), thus removing a likely phosphorylation site. These genes with the point mutation were recombined into pGWB402, a Gateway™ cloning compatible binary vector for expression of these genes, respectively, under control of the CaMV*35S* promoter. These constructs were transformed with *R. rhizogenes* as previously described. One centimeter of each respective root tip was excised and immediately frozen in liquid nitrogen. Frozen tissue was ground and treated with 500 µl of TRIzol™ Reagent (Invitrogen catalog number: 15596026) for five minutes. 100 µl of chloroform was added, the mixture vortexed, and then centrifuged at 13,000 x g at 4°C for 15 min. The aqueous fraction was transferred to a new tube and 250 µl of isopropanol was added and the mixture was incubated at room temperature for 10 min then centrifuged at 13,000 x g at 4°C for 10 min. The supernatant was discarded, and the pellet was rinsed with 1 mL of cold 70% ethanol, then left to dry and rehydrated with 100 µl of nuclease-free H2O. To perform cDNA synthesis, 10 µl of RNA extraction mixture was then added to 1 µl of 10 mM dNTPs (Takara Catalog # 4030) and 100 mM of oligo-dT(18) (R&D Systems Catalog number: RDPC1) primer and incubated at 65°C for 5 min. The final step of cDNA synthesis comprised the addition of 1 µL of M-MLV Reverse Transcriptase (Invitrogen Catalog Number 28025-013), 1 µL RNasin® Ribonuclease Inhibitor (Promega Part # N211A), with 5 µl 5X First Strand Buffer (Invitrogen Catalog Number 18067017) and 6 µl nuclease-free water at 37°C for 60 minutes. Quantitative PCR was performed on a CFX384 Touch Real-Time PCR Detection System (Bio-Rad Laboratories Inc.) using SYBR Green Master Mix as the fluorophore (ThermoFisher Catalog number: 4309155).

These reactions were comprised of 1 µL of cDNA, 3 µL of SYBR Green Master Mix, 0.5 µL of a forward and reverse primer each, and 1 µL of nuclease free water. Four technical replicates were used for each sample, and *SlACT2* (actin; *Solyc11g005330*.*1*) was used as the reference gene for all experiments. Relative expression was calculated using the delta-delta-ct method (Livak and Schmittgen 2001). Transgenic lines of *Solyc04g079980/SlBZR1a* and *Solyc12g089040/BZR1b* were generated by the UC Davis Transformation Facility.

### Quantification of Brassinosteroids

Tomato seeds were surface sterilized and sowed directly on sufficient phosphate MS media (1 mM KH_2_PO_4_) or limiting phosphate (20 µM KH_2_PO_4_) MS media. Shoot and roots were harvested separately at 5 days after germination, slightly dried with kimwipes to remove excess of water and their fresh weight was measured. Approximately 1 gram of tissue per sample was immediately frozen in liquid nitrogen within 3 biological replicates. The samples were subsequently freeze-dried and each weighing cca 5 mg DW was extracted, purified and analyzed using ultra-high performance liquid chromatographic (UHPLC) coupled to electrospray ionization tandem mass spectrometry (ESI–MS/MS) as detailed in (Tarkowská et al., 2016; Ackerman-Lavert et al., 2021).

## Acknowledgements

We thank Tsuyoshi Nakagawa for the pGWB402 plasmid. GSD is supported by the Resnick Sustainability Institute Postdoctoral Fellowship. This work was funded by a BARD Research Grant Agreement No. IS-4827-15 to SMB and SS-G. Additional support was provided by an HHMI Faculty Scholar and NSF 1856749, 2119820 and 1238243 and SS-G has received funding from the European Union’s Horizon 2020 research and innovation program under the Grant Agreement No. [727929] (TOMRES). DT is grateful for financial support from European Regional Development Fund Project, Centre for Experimental Plant Biology No. CZ.02.1.01/0.0/0.0/16_019/0000738.

