## Supplementary information for "The Phosphate Deprivation Response is Mediated by an Interaction between Brassinosteroid Signaling and Zinc in Tomato"

[*] Authors have contributed equally

### Supplementary Figures

**A**

| Source of Variation | % of total variation | P value | P value summary | Significant? | |
|---|---|---|---|---|---|
| Interaction | 1.734 | <0.0001 | **** | Yes | |
| Genotype | 78.50 | <0.0001 | **** | Yes | |
| P treatment | 2.378 | <0.0001 | **** | Yes | |
| ANOVA table | SS (Type III) | DF | MS | F (DFn, DFd) | P value |
| Interaction | 52.75 | 1 | 52.75 | F (1, 185) = 19.20 | P<0.0001 |
| Genotype | 2388 | 1 | 2388 | F (1, 185) = 869.4 | P<0.0001 |
| P treatment | 72.34 | 1 | 72.34 | F (1, 185) = 26.34 | P<0.0001 |
| Residual | 508.1 | 185 | 2.747 | | |

| Tukey's multiple comparisons | Predicted mean diff. | 95.00% CI of diff. | Below threshold? | Adjusted P Value |
|---|---|---|---|---|
| M82:P+ vs. M82:P- | 2.307 | 1.448 to 3.167 | Yes | <0.0001 |
| M82:P+ vs. Penn:P+ | 8.213 | 7.354 to 9.073 | Yes | <0.0001 |
| M82:P+ vs. Penn:P- | 8.395 | 7.477 to 9.313 | Yes | <0.0001 |
| M82:P- vs. Penn:P+ | 5.906 | 5.047 to 6.765 | Yes | <0.0001 |
| M82:P- vs. Penn:P- | 6.088 | 5.170 to 7.006 | Yes | <0.0001 |
| Penn:P+ vs. Penn:P- | 0.1818 | -0.7361 to 1.100 | No | 0.9558 |

**B**

| Source of Variation | % of total variation | P value | P value summary | Significant? | |
|---|---|---|---|---|---|
| Interaction | 5.373 | 0.0010 | ** | Yes | |
| Genotype | 2.642 | 0.0206 | * | Yes | |
| P treatment | 2.057 | 0.0407 | * | Yes | |
| ANOVA table | SS (Type III) | DF | MS | F (DFn, DFd) | P value |
| Interaction | 43.99 | 1 | 43.99 | F (1, 185) = 11.09 | P=0.0010 |
| Genotype | 21.63 | 1 | 21.63 | F (1, 185) = 5.454 | P=0.0206 |
| P treatment | 16.84 | 1 | 16.84 | F (1, 185) = 4.246 | P=0.0407 |
| Residual | 733.7 | 185 | 3.966 | | |

| Tukey's multiple comparisons | Predicted mean diff. | 95.00% CI of diff. | Below threshold? | Adjusted P Value |
|---|---|---|---|---|
| M82:P+ vs. M82:P- | -1.571 | -2.604 to -0.5384 | Yes | 0.0007 |
| M82:P+ vs. Penn:P+ | -0.2900 | -1.323 to 0.7426 | No | 0.8857 |
| M82:P+ vs. Penn:P- | 0.08006 | -1.023 to 1.183 | No | 0.9976 |
| M82:P- vs. Penn:P+ | 1.281 | 0.2484 to 2.314 | Yes | 0.0083 |
| M82:P- vs. Penn:P- | 1.651 | 0.5480 to 2.754 | Yes | 0.0008 |
| Penn:P+ vs. Penn:P- | 0.3701 | -0.7330 to 1.473 | No | 0.8205 |

**C**

| Source of Variation | % of total variation | P value | P value summary | Significant? | |
|---|---|---|---|---|---|
| Interaction | 22.70 | <0.0001 | **** | Yes | |
| Genotype | 62.58 | <0.0001 | **** | Yes | |
| P treatment | 4.871 | <0.0001 | **** | Yes | |
| ANOVA table | SS (Type III) | DF | MS | F (DFn, DFd) | P value |
| Interaction | 1.093 | 1 | 1.093 | F (1, 75) = 169.0 | P<0.0001 |
| Genotype | 3.015 | 1 | 3.015 | F (1, 75) = 466.1 | P<0.0001 |
| P treatment | 0.2347 | 1 | 0.2347 | F (1, 75) = 36.28 | P<0.0001 |
| Residual | 0.4851 | 75 | 0.0065 | | |

| Tukey's multiple comparisons | Predicted mean diff. | 95.00% CI of diff. | Below threshold? | Adjusted P Value |
|---|---|---|---|---|
| M82:P+ vs. M82:P- | -0.1263 | -0.1940 to -0.05862 | Yes | <0.0001 |
| M82:P+ vs. Penn:P+ | -0.6262 | -0.6939 to -0.5585 | Yes | <0.0001 |
| M82:P+ vs. Penn:P- | -0.2818 | -0.3495 to -0.2141 | Yes | <0.0001 |
| M82:P- vs. Penn:P+ | -0.4998 | -0.5667 to -0.4330 | Yes | <0.0001 |
| M82:P- vs. Penn:P- | -0.1554 | -0.2223 to -0.08862 | Yes | <0.0001 |
| Penn:P+ vs. Penn:P- | 0.3444 | 0.2776 to 0.4112 | Yes | <0.0001 |

**D**

| Source of Variation | % of total variation | P value | P value summary | Significant? | |
|---|---|---|---|---|---|
| Interaction | 9.773 | 0.0142 | * | Yes | |
| Genotype | 25.92 | 0.0005 | *** | Yes | |
| P treatment | 50.02 | <0.0001 | **** | Yes | |
| ANOVA table | SS | DF | MS | F (DFn, DFd) | P value |
| Interaction | 4.530 | 1 | 4.530 | F (1, 12) = 8.206 | P=0.0142 |
| Genotype | 12.01 | 1 | 12.01 | F (1, 12) = 21.76 | P=0.0005 |
| P treatment | 23.18 | 1 | 23.18 | F (1, 12) = 42.00 | P<0.0001 |
| Residual | 6.624 | 12 | 0.5520 | | |

| Tukey's multiple comparisons | Mean Diff. | 95.00% CI of diff. | Below threshold? | Adjusted P Value |
|---|---|---|---|---|
| M82:P+ vs. M82:P- | -3.472 | -5.031 to -1.912 | Yes | 0.0001 |
| M82:P+ vs. Penn:P+ | -2.797 | -4.357 to -1.237 | Yes | 0.0009 |
| M82:P+ vs. Penn:P- | -4.140 | -5.700 to -2.581 | Yes | <0.0001 |
| M82:P- vs. Penn:P+ | 0.6746 | -0.8851 to 2.234 | No | 0.5893 |
| M82:P- vs. Penn:P- | -0.6687 | -2.228 to 0.8910 | No | 0.5958 |
| Penn:P+ vs. Penn:P- | -1.343 | -2.903 to 0.2164 | No | 0.1005 |

**Supp. Fig 1: Statistical analysis of Figure 1. A)** Figure 1 Panel A (primary root length). **B)** Figure 1 Panel B (total lateral root length). **C)** Figure 1 Panel D (mature root hair length). **D)** Figure 1 Panel G (pNPP assay).

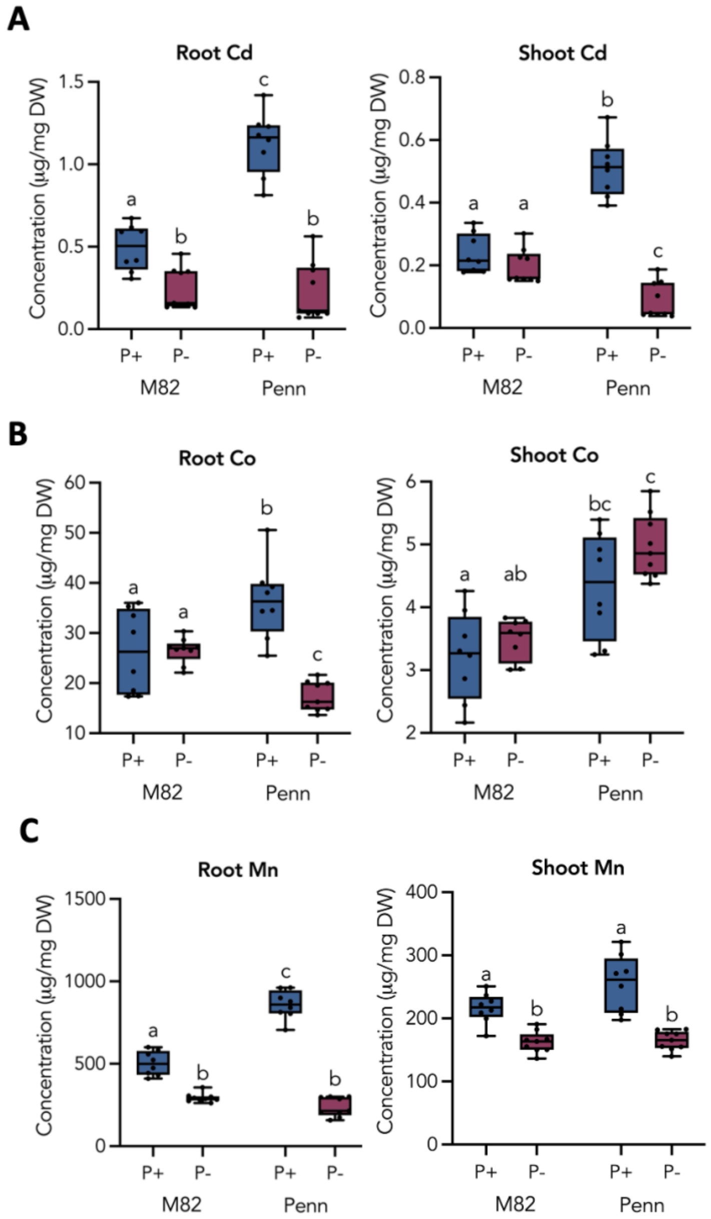

**Supp. Fig. 2: ICP-MS results of Cd, Co, and Mn roots and shoots in P sufficiency or deficiency. A)** Root and shoot Cd **B)** Root and shoot Co and **C)** Root and shoot Mn profile of M82 and Penn in P-sufficient and P-limiting conditions. N= 6 for all plots. Letters represent statistically significant differences as determined by a two-way ANOVA and a post-hoc Tukey test, p<0.01.

**A**

| Source of Variation | % of total variation | P value | P value summary | Significant? | |
|---|---|---|---|---|---|
| Interaction | 0.7111 | 0.2247 | ns | No | |
| Genotype | 1.427 | 0.0894 | ns | No | |
| P treatment | 85.68 | <0.0001 | **** | Yes | |
| ANOVA table | SS (Type III) | DF | MS | F (DFn, DFd) | P value |
| Interaction | 4.420 | 1 | 4.420 | F (1, 29) = 1.539 | P=0.2247 |
| Genotype | 8.869 | 1 | 8.869 | F (1, 29) = 3.089 | P=0.0894 |
| P treatment | 532.5 | 1 | 532.5 | F (1, 29) = 185.5 | P<0.0001 |
| Residual | 83.27 | 29 | 2.871 | | |

| Tukey's multiple comparisons | Predicted mean diff. | 95.00% CI of diff. | Below threshold? | Adjusted P Value |
|---|---|---|---|---|
| M82:P+ vs. M82:P- | 7.341 | 5.098 to 9.585 | Yes | <0.0001 |
| M82:P+ vs. Penn:P+ | -1.778 | -4.168 to 0.6111 | No | 0.2013 |
| M82:P+ vs. Penn:P- | 7.035 | 4.792 to 9.278 | Yes | <0.0001 |
| M82:P- vs. Penn:P+ | -9.120 | -11.45 to -6.793 | Yes | <0.0001 |
| M82:P- vs. Penn:P- | -0.3065 | -2.483 to 1.870 | No | 0.9804 |
| Penn:P+ vs. Penn:P- | 8.813 | 6.487 to 11.14 | Yes | <0.0001 |

**B**

| Source of Variation | % of total variation | P value | P value summary | Significant? | |
|---|---|---|---|---|---|
| Interaction | 1.093 | 0.0259 | * | Yes | |
| Genotype | 3.785 | 0.0001 | *** | Yes | |
| P treatment | 91.39 | <0.0001 | **** | Yes | |
| ANOVA table | SS (Type III) | DF | MS | F (DFn, DFd) | P value |
| Interaction | 6.270 | 1 | 6.270 | F (1, 29) = 5.514 | P=0.0259 |
| Genotype | 21.72 | 1 | 21.72 | F (1, 29) = 19.10 | P=0.0001 |
| P treatment | 524.3 | 1 | 524.3 | F (1, 29) = 461.0 | P<0.0001 |
| Residual | 32.98 | 29 | 1.137 | | |

| Tukey's multiple comparisons | Predicted mean diff. | 95.00% CI of diff. | Below threshold? | Adjusted P Value |
|---|---|---|---|---|
| M82:P+ vs. M82:P- | 7.138 | 5.726 to 8.550 | Yes | <0.0001 |
| M82:P+ vs. Penn:P+ | -2.508 | -4.011 to -1.004 | Yes | 0.0005 |
| M82:P+ vs. Penn:P- | 6.383 | 4.972 to 7.795 | Yes | <0.0001 |
| M82:P- vs. Penn:P+ | -9.646 | -11.11 to -8.181 | Yes | <0.0001 |
| M82:P- vs. Penn:P- | -0.7546 | -2.124 to 0.6150 | No | 0.4498 |
| Penn:P+ vs. Penn:P- | 8.891 | 7.427 to 10.36 | Yes | <0.0001 |

**C**

| Source of Variation | % of total variation | P value | P value summary | Significant? | |
|---|---|---|---|---|---|
| Interaction | 0.1872 | 0.7538 | ns | No | |
| Genotype | 0.4804 | 0.6159 | ns | No | |
| P treatment | 45.60 | <0.0001 | **** | Yes | |
| ANOVA table | SS (Type III) | DF | MS | F (DFn, DFd) | P value |
| Interaction | 0.04914 | 1 | 0.04914 | F (1, 29) = 0.1002 | P=0.7538 |
| Genotype | 0.1261 | 1 | 0.1261 | F (1, 29) = 0.2572 | P=0.6159 |
| P treatment | 11.97 | 1 | 11.97 | F (1, 29) = 24.41 | P<0.0001 |
| Residual | 14.22 | 29 | 0.4903 | | |

| Tukey's multiple comparisons | Predicted mean diff. | 95.00% CI of diff. | Below threshold? | Adjusted P Value |
|---|---|---|---|---|
| M82:P+ vs. M82:P- | -1.133 | -2.060 to -0.2063 | Yes | 0.0120 |
| M82:P+ vs. Penn:P+ | 0.2019 | -0.7855 to 1.189 | No | 0.9439 |
| M82:P+ vs. Penn:P- | -1.087 | -2.014 to -0.1596 | Yes | 0.0168 |
| M82:P- vs. Penn:P+ | 1.335 | 0.3738 to 2.297 | Yes | 0.0038 |
| M82:P- vs. Penn:P- | 0.04670 | -0.8526 to 0.9460 | No | 0.9990 |
| Penn:P+ vs. Penn:P- | -1.288 | -2.250 to -0.3271 | Yes | 0.0053 |

**D**

| Source of Variation | % of total variation | P value | P value summary | Significant? | |
|---|---|---|---|---|---|
| Interaction | 0.3997 | 0.6452 | ns | No | |
| Genotype | 9.799 | 0.0288 | * | Yes | |
| P treatment | 38.05 | <0.0001 | **** | Yes | |
| ANOVA table | SS (Type III) | DF | MS | F (DFn, DFd) | P value |
| Interaction | 0.0005324 | 1 | 0.0005324 | F (1, 28) = 0.2167 | P=0.6452 |
| Genotype | 0.01305 | 1 | 0.01305 | F (1, 28) = 5.312 | P=0.0288 |
| P treatment | 0.05069 | 1 | 0.05069 | F (1, 28) = 20.63 | P<0.0001 |
| Residual | 0.06881 | 28 | 0.002457 | | |

| Tukey's multiple comparisons | Predicted mean diff. | 95.00% CI of diff. | Below threshold? | Adjusted P Value |
|---|---|---|---|---|
| M82:P+ vs. M82:P- | -0.07172 | -0.1375 to -0.005957 | Yes | 0.0286 |
| M82:P+ vs. Penn:P+ | -0.03236 | -0.1024 to 0.03768 | No | 0.5942 |
| M82:P+ vs. Penn:P- | -0.1205 | -0.1881 to -0.05280 | Yes | 0.0002 |
| M82:P- vs. Penn:P+ | 0.03936 | -0.02885 to 0.1076 | No | 0.4083 |
| M82:P- vs. Penn:P- | -0.04875 | -0.1145 to 0.01702 | No | 0.2036 |
| Penn:P+ vs. Penn:P- | -0.08811 | -0.1582 to -0.01806 | Yes | 0.0095 |

**E**

| Source of Variation | % of total variation | P value | P value summary | Significant? | |
|---|---|---|---|---|---|
| Interaction | 48.02 | <0.0001 | **** | Yes | |
| Genotype | 18.37 | 0.0002 | *** | Yes | |
| P treatment | 11.92 | 0.0018 | ** | Yes | |
| ANOVA table | SS (Type III) | DF | MS | F (DFn, DFd) | P value |
| Interaction | 0.8472 | 1 | 0.8472 | F (1, 29) = 47.84 | P<0.0001 |
| Genotype | 0.3241 | 1 | 0.3241 | F (1, 29) = 18.30 | P=0.0002 |
| P treatment | 0.2103 | 1 | 0.2103 | F (1, 29) = 11.87 | P=0.0018 |
| Residual | 0.5136 | 29 | 0.01771 | | |

| Tukey's multiple comparisons | Predicted mean diff. | 95.00% CI of diff. | Below threshold? | Adjusted P Value |
|---|---|---|---|---|
| M82:P+ vs. M82:P- | -0.1617 | -0.3379 to 0.01452 | No | 0.0810 |
| M82:P+ vs. Penn:P+ | -0.5215 | -0.7091 to -0.3338 | Yes | <0.0001 |
| M82:P+ vs. Penn:P- | -0.03875 | -0.2149 to 0.1374 | No | 0.9315 |
| M82:P- vs. Penn:P+ | -0.3598 | -0.5425 to -0.1771 | Yes | <0.0001 |
| M82:P- vs. Penn:P- | 0.1229 | -0.04801 to 0.2938 | No | 0.2266 |
| Penn:P+ vs. Penn:P- | 0.4827 | 0.3000 to 0.6654 | Yes | <0.0001 |

**F**

| Source of Variation | % of total variation | P value | P value summary | Significant? | |
|---|---|---|---|---|---|
| Interaction | 28.51 | <0.0001 | **** | Yes | |
| Genotype | 46.09 | <0.0001 | **** | Yes | |
| P treatment | 22.03 | <0.0001 | **** | Yes | |
| ANOVA table | SS (Type III) | DF | MS | F (DFn, DFd) | P value |
| Interaction | 0.07443 | 1 | 0.07443 | F (1, 29) = 63.02 | P<0.0001 |
| Genotype | 0.1203 | 1 | 0.1203 | F (1, 29) = 101.9 | P<0.0001 |
| P treatment | 0.05750 | 1 | 0.05750 | F (1, 29) = 48.69 | P<0.0001 |
| Residual | 0.03425 | 29 | 0.001181 | | |

| Tukey's multiple comparisons | Predicted mean diff. | 95.00% CI of diff. | Below threshold? | Adjusted P Value |
|---|---|---|---|---|
| M82:P+ vs. M82:P- | -0.01156 | -0.05705 to 0.03394 | No | 0.8993 |
| M82:P+ vs. Penn:P+ | -0.2169 | -0.2654 to -0.1684 | Yes | <0.0001 |
| M82:P+ vs. Penn:P- | -0.03748 | -0.08298 to 0.008019 | No | 0.1353 |
| M82:P- vs. Penn:P+ | -0.2054 | -0.2525 to -0.1582 | Yes | <0.0001 |
| M82:P- vs. Penn:P- | -0.02592 | -0.07006 to 0.01822 | No | 0.3944 |
| Penn:P+ vs. Penn:P- | 0.1794 | 0.1322 to 0.2266 | Yes | <0.0001 |

**Supp. Fig 3: Statistical analysis of Figure 3. A)** Figure 3 Panel A (Root P). **B)** Figure 3 Panel B (Shoot P). **C)** Figure 3 Panel C (Root Fe). **D)** Figure 3 Panel D (Shoot Fe). **E)** Figure 3 Panel E (Root Zn). **F)** Figure 3 Panel F (Shoot Zn).

**A**

| Source of Variation | % of total variation | P value | P value summary | Significant? | |
|---|---|---|---|---|---|
| Interaction | 0.7683 | 0.2444 | ns | No | |
| Zn treatment | 0.8349 | 0.2251 | ns | No | |
| P treatment | 34.88 | <0.0001 | **** | Yes | |
| ANOVA table | SS (Type III) | DF | MS | F (DFn, DFd) | P value |
| Interaction | 5.141 | 1 | 5.141 | F (1, 113) = 1.369 | P=0.2444 |
| Zn treatment | 5.586 | 1 | 5.586 | F (1, 113) = 1.488 | P=0.2251 |
| P treatment | 233.3 | 1 | 233.3 | F (1, 113) = 62.15 | P<0.0001 |
| Residual | 424.3 | 113 | 3.755 | | |

| Tukey's multiple comparisons | Predicted mean diff. | 95.00% CI of diff. | Below threshold? | Adjusted P Value |
|---|---|---|---|---|
| M82 Zn+:P+ vs. M82 Zn+:P- | 2.410 | 1.080 to 3.740 | Yes | <0.0001 |
| M82 Zn+:P+ vs. M82 Zn-:P+ | -0.8578 | -2.141 to 0.4256 | No | 0.3066 |
| M82 Zn+:P+ vs. M82 Zn-:P- | 2.392 | 1.075 to 3.709 | Yes | <0.0001 |
| M82 Zn+:P- vs. M82 Zn-:P+ | -3.268 | -4.598 to -1.937 | Yes | <0.0001 |
| M82 Zn+:P- vs. M82 Zn-:P- | -0.01780 | -1.381 to 1.345 | No | >0.9999 |
| M82 Zn-:P+ vs. M82 Zn-:P- | 3.250 | 1.932 to 4.567 | Yes | <0.0001 |

| Source of Variation | % of total variation | P value | P value summary | Significant? | |
|---|---|---|---|---|---|
| Interaction | 0.4863 | 0.4343 | ns | No | |
| Zn treatment | 0.8453 | 0.3031 | ns | No | |
| P treatment | 0.6541 | 0.3648 | ns | No | |
| ANOVA table | SS (Type III) | DF | MS | F (DFn, DFd) | P value |
| Interaction | 0.8244 | 1 | 0.8244 | F (1, 124) = 0.6152 | P=0.4343 |
| Zn treatment | 1.433 | 1 | 1.433 | F (1, 124) = 1.069 | P=0.3031 |
| P treatment | 1.109 | 1 | 1.109 | F (1, 124) = 0.8275 | P=0.3648 |
| Residual | 166.2 | 124 | 1.340 | | |

| Tukey's multiple comparisons | Predicted mean diff. | 95.00% CI of diff. | Below threshold? | Adjusted P Value |
|---|---|---|---|---|
| Penn Zn+:P+ vs. Penn Zn+:P- | -0.3481 | -1.127 to 0.4307 | No | 0.6507 |
| Penn Zn+:P+ vs. Penn Zn-:P+ | -0.3737 | -1.139 to 0.3921 | No | 0.5832 |
| Penn Zn+:P+ vs. Penn Zn-:P- | -0.3994 | -1.133 to 0.3346 | No | 0.4912 |
| Penn Zn+:P- vs. Penn Zn-:P+ | -0.02557 | -0.8044 to 0.7533 | No | 0.9998 |
| Penn Zn+:P- vs. Penn Zn-:P- | -0.05133 | -0.7990 to 0.6964 | No | 0.9980 |
| Penn Zn-:P+ vs. Penn Zn-:P- | -0.02575 | -0.7598 to 0.7083 | No | 0.9997 |

**B**

| Source of Variation | % of total variation | P value | P value summary | Significant? | |
|---|---|---|---|---|---|
| Interaction | 0.02565 | 0.8503 | ns | No | |
| Genotype | 0.6825 | 0.3314 | ns | No | |
| P treatment | 28.53 | <0.0001 | **** | Yes | |
| ANOVA table | SS (Type III) | DF | MS | F (DFn, DFd) | P value |
| Interaction | 0.04753 | 1 | 0.04753 | F (1, 98) = 0.03580 | P=0.8503 |
| Zn treatment | 1.265 | 1 | 1.265 | F (1, 98) = 0.9527 | P=0.3314 |
| P treatment | 52.86 | 1 | 52.86 | F (1, 98) = 39.82 | P<0.0001 |
| Residual | 130.1 | 98 | 1.327 | | |

| Tukey's multiple comparisons | Predicted mean diff. | 95.00% CI of diff. | Below threshold? | Adjusted P Value |
|---|---|---|---|---|
| M82 Zn+:P+ vs. M82 Zn+:P- | -1.493 | -2.372 to -0.6146 | Yes | 0.0001 |
| M82 Zn+:P+ vs. M82 Zn-:P+ | -0.2677 | -1.146 to 0.6110 | No | 0.8559 |
| M82 Zn+:P+ vs. M82 Zn-:P- | -1.674 | -2.503 to -0.8454 | Yes | <0.0001 |
| M82 Zn+:P- vs. M82 Zn-:P+ | 1.226 | 0.3563 to 2.095 | Yes | 0.0021 |
| M82 Zn+:P- vs. M82 Zn-:P- | -0.1808 | -0.9995 to 0.6380 | No | 0.9387 |
| M82 Zn-:P+ vs. M82 Zn-:P- | -1.406 | -2.225 to -0.5876 | Yes | 0.0001 |

| Source of Variation | % of total variation | P value | P value summary | Significant? | |
|---|---|---|---|---|---|
| Interaction | 9.415 | <0.0001 | **** | Yes | |
| Genotype | 12.17 | <0.0001 | **** | Yes | |
| P treatment | 0.1294 | 0.6351 | ns | No | |
| ANOVA table | SS (Type III) | DF | MS | F (DFn, DFd) | P value |
| Interaction | 15.83 | 1 | 15.83 | F (1, 133) = 16.47 | P<0.0001 |
| Zn treatment | 20.46 | 1 | 20.46 | F (1, 133) = 21.29 | P<0.0001 |
| P treatment | 0.2175 | 1 | 0.2175 | F (1, 133) = 0.2263 | P=0.6351 |
| Residual | 127.8 | 133 | 0.9612 | | |

| Tukey's multiple comparisons | Predicted mean diff. | 95.00% CI of diff. | Below threshold? | Adjusted P Value |
|---|---|---|---|---|
| Penn Zn+:P+ vs. Penn Zn+:P- | 0.7684 | 0.1754 to 1.361 | Yes | 0.0054 |
| Penn Zn+:P+ vs. Penn Zn-:P+ | -0.09424 | -0.7470 to 0.5585 | No | 0.9819 |
| Penn Zn+:P+ vs. Penn Zn-:P- | -0.7014 | -1.294 to -0.1084 | Yes | 0.0134 |
| Penn Zn+:P- vs. Penn Zn-:P+ | -0.8627 | -1.515 to -0.2099 | Yes | 0.0043 |
| Penn Zn+:P- vs. Penn Zn-:P- | -1.470 | -2.063 to -0.8768 | Yes | <0.0001 |
| Penn Zn-:P+ vs. Penn Zn-:P- | -0.6072 | -1.260 to 0.04560 | No | 0.0782 |

**C**

| Source of Variation | % of total variation | P value | P value summary | Significant? | |
|---|---|---|---|---|---|
| Zn treatment | 0.2871 | 0.6877 | ns | No | |
| P treatment | 20.80 | 0.0013 | ** | Yes | |
| ANOVA table | SS (Type III) | DF | MS | F (DFn, DFd) | P value |
| Zn treatment | 0.001562 | 1 | 0.001562 | F (1, 45) = 0.1637 | P=0.6877 |
| P treatment | 0.1132 | 1 | 0.1132 | F (1, 45) = 11.86 | P=0.0013 |
| Residual | 0.4294 | 45 | 0.009542 | | |

| Source of Variation | % of total variation | P value | P value summary | Significant? | |
|---|---|---|---|---|---|
| Zn treatment | 13.10 | 0.0104 | * | Yes | |
| P treatment | 6.902 | 0.0585 | ns | No | |
| ANOVA table | SS (Type III) | DF | MS | F (DFn, DFd) | P value |
| Zn treatment | 0.08090 | 1 | 0.08090 | F (1, 44) = 7.164 | P=0.0104 |
| P treatment | 0.04262 | 1 | 0.04262 | F (1, 44) = 3.774 | P=0.0585 |
| Residual | 0.4968 | 44 | 0.01129 | | |

| Tukey's multiple comparisons | Predicted mean diff. | 95.00% CI of diff. | Below threshold? | Adjusted P Value |
|---|---|---|---|---|
| Zn+:P+ vs. Zn+:P- | -0.09712 | -0.1723 to -0.02190 | Yes | 0.0066 |
| Zn+:P+ vs. Zn-:P+ | 0.01141 | -0.06382 to 0.08664 | No | 0.9773 |
| Zn+:P+ vs. Zn-:P- | -0.08571 | -0.1921 to 0.02067 | No | 0.1534 |
| Zn+:P- vs. Zn-:P+ | 0.1085 | 0.002148 to 0.2149 | Yes | 0.0440 |
| Zn+:P- vs. Zn-:P- | 0.01141 | -0.06382 to 0.08664 | No | 0.9773 |
| Zn-:P+ vs. Zn-:P- | -0.09712 | -0.1723 to -0.02190 | Yes | 0.0066 |

| Tukey's multiple comparisons | Predicted mean diff. | 95.00% CI of diff. | Below threshold? | Adjusted P Value |
|---|---|---|---|---|
| Zn+:P+ vs. Zn+:P- | 0.06037 | -0.02260 to 0.1433 | No | 0.2255 |
| Zn+:P+ vs. Zn-:P+ | 0.08317 | 0.0002053 to 0.1661 | Yes | 0.0492 |
| Zn+:P+ vs. Zn-:P- | 0.1435 | 0.02472 to 0.2624 | Yes | 0.0122 |
| Zn+:P- vs. Zn-:P+ | 0.02280 | -0.09302 to 0.1386 | No | 0.9524 |
| Zn+:P- vs. Zn-:P- | 0.08317 | 0.0002053 to 0.1661 | Yes | 0.0492 |
| Zn-:P+ vs. Zn-:P- | 0.06037 | -0.02260 to 0.1433 | No | 0.2255 |

**Supp. Fig 4: Statistical analysis of Fig 4. A)** Primary root responses of M82 (left) and *S. pennellii* (right) **B)** Total lateral root length responses of M82 (left) and *S. pennellii* (right) **C)** Mature root hair length responses of M82 (left) and *S. pennellii* (right) in P sufficiency or deficiency when Zn is absent or present.

| Derivative | Source | p.value | adjusted_p.value |
|---|---|---|---|
| CR | P treatment | 0.9029 | 0.9612 |
| CN | P treatment | 0.4129 | 0.7570 |
| 6-deoxoCT | P treatment | 0.9786 | 0.9786 |
| 6-deoxoTY | P treatment | 0.2105 | 0.4961 |
| 6-oxoCN | P treatment | 0.2634 | 0.5433 |
| CS | P treatment | 0.5973 | 0.7884 |
| BL | P treatment | 0.7968 | 0.9067 |
| homoCS | P treatment | 0.2421 | 0.5326 |
| norTE | P treatment | 0.0068 | 0.0448 |
| norCS | P treatment | 0.1250 | 0.4126 |
| homoBL | P treatment | 0.2010 | 0.4961 |
| CR | Species | 0.0003 | 0.0035 |
| CN | Species | 0.0199 | 0.0937 |
| 6-deoxoCT | Species | 0.0772 | 0.3186 |
| 6-deoxoTY | Species | 0.0001 | 0.0020 |
| 6-oxoCN | Species | 0.4706 | 0.7727 |
| CS | Species | 0.1433 | 0.4298 |
| BL | Species | 0.0056 | 0.0448 |
| homoCS | Species | 0.6732 | 0.8545 |
| norTE | Species | 0.5475 | 0.7727 |
| norCS | Species | 0.0000 | 0.0009 |
| homoBL | Species | 0.4955 | 0.7727 |
| CR | Interaction | 0.4771 | 0.7727 |
| CN | Interaction | 0.3277 | 0.6361 |
| 6-deoxoCT | Interaction | 0.1583 | 0.4354 |
| 6-deoxoTY | Interaction | 0.7339 | 0.8764 |
| 6-oxoCN | Interaction | 0.8375 | 0.9212 |
| CS | Interaction | 0.0138 | 0.0760 |
| BL | Interaction | 0.5439 | 0.7727 |
| homoCS | Interaction | 0.7436 | 0.8764 |
| norTE | Interaction | 0.1182 | 0.4126 |
| norCS | Interaction | 0.9737 | 0.9786 |
| homoBL | Interaction | 0.5619 | 0.7727 |

**Supp. Fig 5: Statistical analysis of BR levels in M82 and *S. pennellii* (Figure 5C).** P-values are obtained from a two-way ANOVA analysis. BL is highlighted as one of my main types of BRs. The samples were clustered as follows: z-scores were calculated for each BR level, Euclidean distances between samples were calculated and hierarchical clustering was performed using the complete method. Hypothesis testing: Two-way ANOVA was performed for each BR with P and Species as main factors and their interaction. For each BR with significant effect, post hoc comparisons were performed using Tukey's HSD test. Fig. 5A,B For each species one way analysis of variance was performed with hormone concentration as the independent variable. When significant the ANOVA was followed by Tukey's HSD test to compare between the different concentration pairs. Fig 5 D,E - For M82, two independent experiments were analyzed. Thus, to determine the effect of P and BRZ on LR, mixed model analysis of variance (ANOVA) was performed with P, BRZ and their interaction as fixed factors and the batch as random factor. Post-hoc analysis was performed using pairwise differences of LS-mean as implemented in the lmerTest R package. P-values are based on the t-distribution using degrees of freedom according to Satterthwaites method. P-values that passed correction for multiple comparisons using the False Discovery Rate procedure with alpha<0 are indicated by letters. For Penn two-way ANOVA was performed with P and BRZ treatments as main factors and their interaction.

**A**

M82 BL p values

| | diff | lwr | upr | p adj |
|---|---|---|---|---|
| 0.01-0 | -0.1188333 | -2.147240 | 1.9095738 | 0.9997857 |
| 0.1-0 | -0.2318333 | -2.095492 | 1.6318256 | 0.9958823 |
| 1-0 | -0.1673333 | -2.101344 | 1.7666769 | 0.9989998 |
| 10-0 | -1.7234333 | -3.751840 | 0.3049738 | 0.1232693 |
| 0.1-0.01 | -0.1130000 | -2.074444 | 1.8484444 | 0.9997995 |
| 1-0.01 | -0.0485000 | -2.076907 | 1.9799071 | 0.9999940 |
| 10-0.01 | -1.6046000 | -3.723202 | 0.5140021 | 0.2025182 |
| 1-0.1 | 0.0645000 | -1.799159 | 1.9281589 | 0.9999736 |
| 10-0.1 | -1.4916000 | -3.453044 | 0.4698444 | 0.1993606 |
| 10-1 | -1.5561000 | -3.584507 | 0.4723071 | 0.1926136 |

Penn BL p values

| | diff | lwr | upr | p adj |
|---|---|---|---|---|
| 0.01-0 | 0.77875000 | -0.2395590 | 1.7970590 | 0.2078131 |
| 0.1-0 | -1.83175000 | -2.9350599 | -0.7284401 | 0.0002297 |
| 1-0 | -1.74339286 | -2.9472027 | -0.5395830 | 0.0015024 |
| 10-0 | -1.61337500 | -2.7763658 | -0.4503842 | 0.0025271 |
| 0.1-0.01 | -2.61050000 | -3.5600780 | -1.6609220 | 0.0000000 |
| 1-0.01 | -2.52214286 | -3.5868323 | -1.4574534 | 0.0000003 |
| 10-0.01 | -2.39212500 | -3.4104340 | -1.3738160 | 0.0000004 |
| 1-0.1 | 0.08835714 | -1.0578991 | 1.2346134 | 0.9994635 |
| 10-0.1 | 0.21837500 | -0.8849349 | 1.3216849 | 0.9796175 |
| 10-1 | 0.13001786 | -1.0737920 | 1.3338277 | 0.9979800 |

**B**

M82 BRZ p values

| | diff | lwr | upr | p adj |
|---|---|---|---|---|
| 0.003-0 | 2.0091667 | 0.1756713 | 3.8426620 | 0.0271323 |
| 0.03-0 | -0.4476333 | -2.1676044 | 1.2723377 | 0.9359405 |
| 0.3-0 | -0.1285000 | -1.7684281 | 1.5114281 | 0.9992991 |
| 3-0 | -3.5875000 | -5.2274281 | -1.9475719 | 0.0000144 |
| 0.03-0.003 | -2.4568000 | -4.3622242 | -0.5513758 | 0.0073857 |
| 0.3-0.003 | -2.1376667 | -3.9711620 | -0.3041713 | 0.0170820 |
| 3-0.003 | -5.5966667 | -7.4301620 | -3.7631713 | 0.0000001 |
| 0.3-0.03 | 0.3191333 | -1.4008377 | 2.0391044 | 0.9807287 |
| 3-0.03 | -3.1398667 | -4.8598377 | -1.4198956 | 0.0001718 |
| 3-0.3 | -3.4590000 | -5.0989281 | -1.8190719 | 0.0000244 |

Penn BRZ p values

| | diff | lwr | upr | p adj |
|---|---|---|---|---|
| 0.003-0 | 0.53619444 | -0.5375301 | 1.6099190 | 0.6216528 |
| 0.03-0 | 0.09305000 | -0.9551060 | 1.1412060 | 0.9990875 |
| 0.3-0 | 0.05008333 | -0.9585053 | 1.0586719 | 0.9999087 |
| 3-0 | 0.01521667 | -0.9521875 | 0.9826209 | 0.9999991 |
| 0.03-0.003 | -0.44314444 | -1.4584347 | 0.5721458 | 0.7305423 |
| 0.3-0.003 | -0.48611111 | -1.4605003 | 0.4882780 | 0.6225262 |
| 3-0.003 | -0.52097778 | -1.4526720 | 0.4107165 | 0.5148184 |
| 0.3-0.03 | -0.04296667 | -0.9891066 | 0.9031733 | 0.9999360 |
| 3-0.03 | -0.07783333 | -0.9799424 | 0.8242757 | 0.9991844 |
| 3-0.3 | -0.03486667 | -0.8906825 | 0.8209491 | 0.9999585 |

**C**

| Source of Variation | % of total variation | P value | P value summary | Significant? |
|---|---|---|---|---|
| Interaction | 18.80 | <0.0001 | **** | Yes |
| Genotype | 22.05 | <0.0001 | **** | Yes |
| BRZ treatment | 13.44 | <0.0001 | **** | Yes |

| ANOVA table | SS (Type III) | DF | MS | F (DFn, DFd) | P value |
|---|---|---|---|---|---|
| Interaction | 221.5 | 3 | 73.83 | F (3, 95) = 12.67 | P<0.0001 |
| Genotype | 259.8 | 1 | 259.8 | F (1, 95) = 44.58 | P<0.0001 |
| BRZ treatment | 158.3 | 3 | 52.78 | F (3, 95) = 9.058 | P<0.0001 |
| Residual | 553.5 | 95 | 5.827 | | |

**D**

| Source of Variation | % of total variation | P value | P value summary | Significant? |
|---|---|---|---|---|
| Interaction | 10.18 | 0.0001 | *** | Yes |
| Genotype | 7.896 | <0.0001 | **** | Yes |
| BRZ treatment | 9.508 | 0.0002 | *** | Yes |

| ANOVA table | SS (Type III) | DF | MS | F (DFn, DFd) | P value |
|---|---|---|---|---|---|
| Interaction | 183.4 | 3 | 61.13 | F (3, 143) = 7.478 | P=0.0001 |
| Genotype | 142.3 | 1 | 142.3 | F (1, 143) = 17.40 | P<0.0001 |
| BRZ treatment | 171.3 | 3 | 57.11 | F (3, 143) = 6.985 | P=0.0002 |
| Residual | 1169 | 143 | 8.176 | | |

**Supp. Fig 6: Statistical analysis of Figure 5 for A)** M82 and Penn primary root length measurement upon BL and **B)** BRZ treatment. **C)** LR number and **D)** LR length of mock and BRZ-treated (3 µM) M82 and Penn plants.

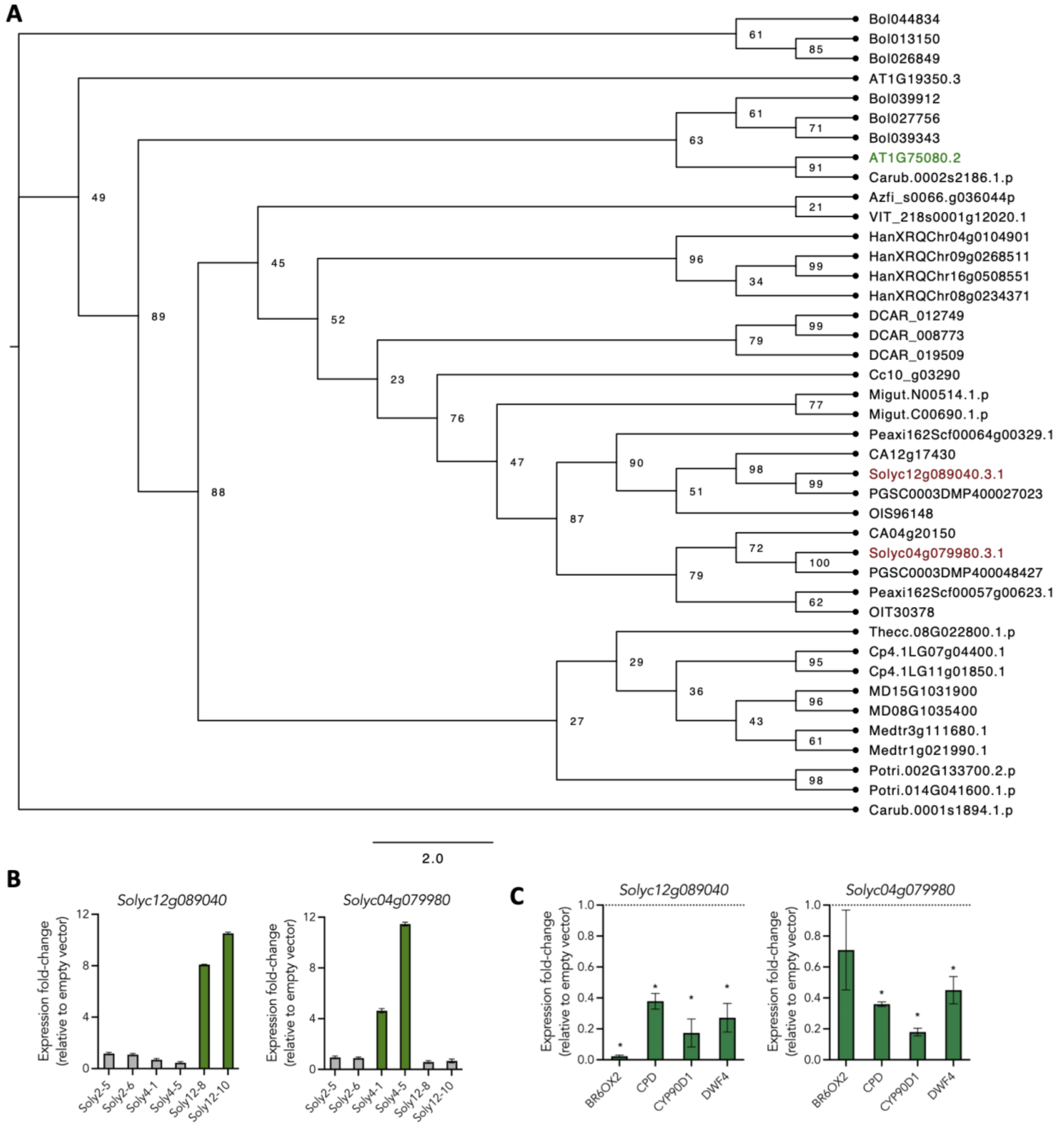

**Supp. Fig 7: BZR1/BES1 phylogeny in tomato. A)** *Solyc12g089040* and *Solyc04g079980* genes were mutated in tomato hairy root lines. **B)** qPCR validation of overexpression of *Solyc12g089040* and *Solyc04g079980* in mutant hairy root lines. Soly12-8 and Soly12-10 lines showing promising results for *Solyc12g089040,* and Soly4-1 and Soly4-5 showing promising results for *Solyc04g079980.* **C)** Soly12-10 and Soly4-5 lines were used to check expression of downstream BZR1 target genes using qPCR, and stable mutants of these lines were generated.

**A**

| Source of Variation | % of total variation | P value | P value summary | Significant? | |
|---|---|---|---|---|---|
| Interaction | 7.171 | 0.0003 | *** | Yes | |
| Genotype | 13.61 | <0.0001 | **** | Yes | |
| BRZ treatment | 65.16 | <0.0001 | **** | Yes | |
| ANOVA table | SS (Type III) | DF | MS | F (DFn, DFd) | P value |
| Interaction | 27.32 | 1 | 27.32 | F (1, 53) = 14.87 | P=0.0003 |
| Genotype | 51.85 | 1 | 51.85 | F (1, 53) = 28.22 | P<0.0001 |
| BRZ treatment | 248.3 | 1 | 248.3 | F (1, 53) = 135.1 | P<0.0001 |
| Residual | 97.37 | 53 | 1.837 | | |

| Tukey's multiple comparisons | Predicted mean diff. | 95.00% CI of diff. | Below threshold? | Adjusted P Value |
|---|---|---|---|---|
| WT:BRZ- vs. WT:BRZ+ | 5.698 | 4.186 to 7.210 | Yes | <0.0001 |
| WT:BRZ- vs. SlBZR1-D:BRZ- | -0.5360 | -1.878 to 0.8064 | No | 0.7156 |
| WT:BRZ- vs. SlBZR1-D:BRZ+ | 2.323 | 1.015 to 3.632 | Yes | 0.0001 |
| WT:BRZ+ vs. SlBZR1-D:BRZ- | -6.234 | -7.683 to -4.785 | Yes | <0.0001 |
| WT:BRZ+ vs. SlBZR1-D:BRZ+ | -3.375 | -4.793 to -1.957 | Yes | <0.0001 |
| SlBZR1-D:BRZ- vs. SlBZR1-D:BRZ+ | 2.859 | 1.624 to 4.095 | Yes | <0.0001 |

**B**

| Source of Variation | % of total variation | P value | P value summary | Significant? | |
|---|---|---|---|---|---|
| Interaction | 16.02 | <0.0001 | **** | Yes | |
| Genotype | 3.437 | 0.0039 | ** | Yes | |
| P treatment | 22.50 | <0.0001 | **** | Yes | |
| ANOVA table | SS (Type III) | DF | MS | F (DFn, DFd) | P value |
| Interaction | 119.9 | 1 | 119.9 | F (1, 147) = 40.09 | P<0.0001 |
| Genotype | 25.72 | 1 | 25.72 | F (1, 147) = 8.598 | P=0.0039 |
| P treatment | 168.4 | 1 | 168.4 | F (1, 147) = 56.29 | P<0.0001 |
| Residual | 439.8 | 147 | 2.992 | | |

| Tukey's multiple comparisons | Predicted mean diff. | 95.00% CI of diff. | Below threshold? | Adjusted P Value |
|---|---|---|---|---|
| wt:P+ vs. wt:P- | 3.895 | 2.850 to 4.940 | Yes | <0.0001 |
| wt:P+ vs. bzr1:P+ | 0.9572 | -0.07431 to 1.989 | No | 0.0793 |
| wt:P+ vs. bzr1:P- | 1.287 | 0.2489 to 2.325 | Yes | 0.0084 |
| wt:P- vs. bzr1:P+ | -2.938 | -3.970 to -1.907 | Yes | <0.0001 |
| wt:P- vs. bzr1:P- | -2.609 | -3.647 to -1.570 | Yes | <0.0001 |
| bzr1:P+ vs. bzr1:P- | 0.3298 | -0.6947 to 1.354 | No | 0.8370 |

**C**

| Source of Variation | % of total variation | P value | P value summary | Significant? | |
|---|---|---|---|---|---|
| Interaction | 7.722 | 0.0001 | *** | Yes | |
| Genotype | 2.972 | 0.0146 | * | Yes | |
| P treatment | 17.43 | <0.0001 | **** | Yes | |
| ANOVA table | SS (Type III) | DF | MS | F (DFn, DFd) | P value |
| Interaction | 265.9 | 1 | 265.9 | F (1, 149) = 15.85 | 265.9 |
| Genotype | 102.3 | 1 | 102.3 | F (1, 149) = 6.101 | 102.3 |
| P treatment | 600.0 | 1 | 600.0 | F (1, 149) = 35.78 | 600.0 |
| Residual | 2499 | 149 | 16.77 | | 2499 |

| Tukey's multiple comparisons | Predicted mean diff. | 95.00% CI of diff. | Below threshold? | Adjusted P Value |
|---|---|---|---|---|
| wt:P+ vs. wt:P- | -6.601 | -9.074 to -4.127 | Yes | <0.0001 |
| wt:P+ vs. bzr1:P+ | -1.002 | -3.428 to 1.425 | No | 0.7070 |
| wt:P+ vs. bzr1:P- | -2.326 | -4.768 to 0.1153 | No | 0.0680 |
| wt:P- vs. bzr1:P+ | 5.599 | 3.172 to 8.026 | Yes | <0.0001 |
| wt:P- vs. bzr1:P- | 4.274 | 1.832 to 6.716 | Yes | <0.0001 |
| bzr1:P+ vs. bzr1:P- | -1.325 | -3.719 to 1.069 | No | 0.4779 |

**D**

| Source of Variation | % of total variation | P value | P value summary | Significant? | |
|---|---|---|---|---|---|
| Interaction | 65.82 | <0.0001 | **** | Yes | |
| Genotype | 7.008 | <0.0001 | **** | Yes | |
| P treatment | 2.014 | <0.0001 | **** | Yes | |
| ANOVA table | SS (Type III) | DF | MS | F (DFn, DFd) | P value |
| Interaction | 63.12 | 1 | 63.12 | F (1, 779) = 2002 | P<0.0001 |
| Genotype | 6.720 | 1 | 6.720 | F (1, 779) = 213.2 | P<0.0001 |
| P treatment | 1.931 | 1 | 1.931 | F (1, 779) = 61.24 | P<0.0001 |
| Residual | 24.56 | 779 | 0.03153 | | |

| Tukey's multiple comparisons | Predicted mean diff. | 95.00% CI of diff. | Below threshold? | Adjusted P Value |
|---|---|---|---|---|
| M82:P+ vs. M82:P- | -0.6808 | -0.7280 to -0.6337 | Yes | <0.0001 |
| M82:P+ vs. bzr1:P+ | -0.7686 | -0.8169 to -0.7202 | Yes | <0.0001 |
| M82:P+ vs. bzr1:P- | -0.2904 | -0.3328 to -0.2481 | Yes | <0.0001 |
| M82:P- vs. bzr1:P+ | -0.08773 | -0.1392 to -0.0362 | Yes | <0.0001 |
| M82:P- vs. bzr1:P- | 0.3904 | 0.3445 to 0.4363 | Yes | <0.0001 |
| bzr1:P+ vs. bzr1:P- | 0.4781 | 0.4309 to 0.5253 | Yes | <0.0001 |

**E**

| Source of Variation | % of total variation | P value | P value summary | Significant? | |
|---|---|---|---|---|---|
| Interaction | 6.276 | <0.0001 | **** | Yes | |
| Genotype | 19.91 | <0.0001 | **** | Yes | |
| P treatment | 69.91 | <0.0001 | **** | Yes | |
| ANOVA table | SS | DF | MS | F (DFn, DFd) | P value |
| Interaction | 7416744 | 1 | 7416744 | F (1, 96) = 154.4 | P<0.0001 |
| Genotype | 23531910 | 1 | 23531910 | F (1, 96) = 489.8 | P<0.0001 |
| P treatment | 82621374 | 1 | 82621374 | F (1, 96) = 1720 | P<0.0001 |
| Residual | 4612413 | 96 | 48046 | | |

| Tukey's multiple comparisons | Predicted mean diff. | 95.00% CI of diff. | Below threshold? | Adjusted P Value |
|---|---|---|---|---|
| M82:P+ vs. M82:P- | -1273 | -1435 to -1111 | Yes | <0.0001 |
| M82:P+ vs. bzr1:P+ | -425.5 | -587.6 to -263.4 | Yes | <0.0001 |
| M82:P+ vs. bzr1:P- | -2788 | -2950 to -2626 | Yes | <0.0001 |
| M82:P- vs. bzr1:P+ | 847.7 | 685.6 to 1010 | Yes | <0.0001 |
| M82:P- vs. bzr1:P- | -1515 | -1677 to -1353 | Yes | <0.0001 |
| bzr1:P+ vs. bzr1:P- | -2363 | -2525 to -2201 | Yes | <0.0001 |

**Supp. Fig 8: Statistical analysis of Figure 6. A)** Figure 6 Panel A (hypocotyl length). **B)** Figure 6 Panel C (primary root length). **C)** Figure 6 Panel D (total lateral root length). **D)** Figure 6 Panel E (mature root hair length). **E)** Figure 6 Panel F (BCIP staining intensity).

**A**

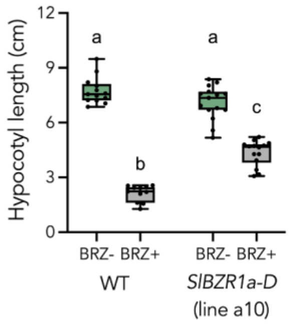

**B**

| Source of Variation | % of total variation | P value | P value summary | Significant? |
|---|---|---|---|---|
| Interaction | 9.768 | <0.0001 | **** | Yes |
| Genotype | 2.977 | 0.0003 | *** | Yes |
| BRZ treatment | 83.39 | <0.0001 | **** | Yes |
| ANOVA table | SS (Type III) | DF | MS | F (DFn, DFd) |
| Interaction | 26.75 | 1 | 26.75 | F (1, 49) = 48.79 |
| Genotype | 8.153 | 1 | 8.153 | F (1, 49) = 14.87 |
| BRZ treatment | 228.4 | 1 | 228.4 | F (1, 49) = 416.5 |
| Residual | 26.87 | 49 | 0.5483 | |

| Tukey's multiple comparisons | Predicted mean diff. | 95.00% CI of diff. | Below threshold? | Adjusted P Value |
|---|---|---|---|---|
| WT:BRZ- vs. WT:BRZ+ | 5.610 | 4.803 to 6.417 | Yes | <0.0001 |
| WT:BRZ- vs. SlBZR1-D:BRZ- | 0.6407 | -0.1055 to 1.387 | No | 0.1159 |
| WT:BRZ- vs. SlBZR1-D:BRZ+ | 3.390 | 2.631 to 4.148 | Yes | <0.0001 |
| WT:BRZ+ vs. SlBZR1-D:BRZ- | -4.969 | -5.751 to -4.187 | Yes | <0.0001 |
| WT:BRZ+ vs. SlBZR1-D:BRZ+ | -2.220 | -3.013 to -1.427 | Yes | <0.0001 |
| SlBZR1-D:BRZ- vs. SlBZR1-D:BRZ+ | 2.749 | 2.017 to 3.481 | Yes | <0.0001 |

**C**

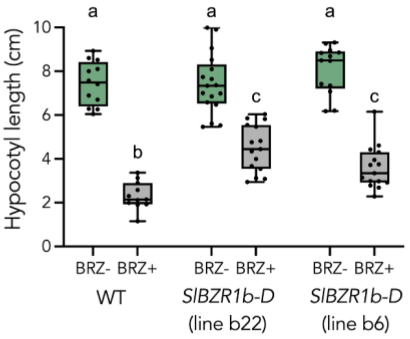

**D**

| Source of Variation | % of total variation | P value | P value summary | Significant? |
|---|---|---|---|---|
| Interaction | 3.889 | 0.0009 | *** | Yes |
| Genotype | 3.781 | 0.0011 | ** | Yes |
| BRZ treatment | 76.38 | <0.0001 | **** | Yes |
| ANOVA table | SS (Type III) | DF | MS | F (DFn, DFd) |
| Interaction | 18.02 | 2 | 9.009 | F (2, 77) = 7.721 |
| Genotype | 17.52 | 2 | 8.759 | F (2, 77) = 7.507 |
| BRZ treatment | 353.9 | 1 | 353.9 | F (1, 77) = 303.3 |
| Residual | 89.84 | 77 | 1.167 | |

| Tukey's multiple comparisons | Predicted mean diff. | 95.00% CI of diff. | Below threshold? | Adjusted P Value |
|---|---|---|---|---|
| WT:BRZ- vs. WT:BRZ+ | 5.123 | 3.805 to 6.440 | Yes | <0.0001 |
| WT:BRZ- vs. SlBZR1b-D b22:BRZ- | 0.02487 | -1.165 to 1.215 | No | >0.9999 |
| WT:BRZ- vs. SlBZR1b-D b22:BRZ+ | 2.947 | 1.725 to 4.170 | Yes | <0.0001 |
| WT:BRZ- vs. SlBZR1b-D b6:BRZ- | -0.6402 | -1.904 to 0.6236 | No | 0.6776 |
| WT:BRZ- vs. SlBZR1b-D b6:BRZ+ | 3.840 | 2.617 to 5.062 | Yes | <0.0001 |
| WT:BRZ+ vs. SlBZR1b-D b22:BRZ- | -5.098 | -6.319 to -3.876 | Yes | <0.0001 |
| WT:BRZ+ vs. SlBZR1b-D b22:BRZ+ | -2.175 | -3.428 to -0.9220 | Yes | <0.0001 |
| WT:BRZ+ vs. SlBZR1b-D b6:BRZ- | -5.763 | -7.056 to -4.469 | Yes | <0.0001 |
| WT:BRZ+ vs. SlBZR1b-D b6:BRZ+ | -1.283 | -2.536 to -0.02965 | Yes | 0.0417 |
| SlBZR1b-D b22:BRZ- vs. SlBZR1b-D b22:BRZ+ | 2.923 | 1.804 to 4.041 | Yes | <0.0001 |
| SlBZR1b-D b22:BRZ- vs. SlBZR1b-D b6:BRZ- | -0.6651 | -1.828 to 0.4981 | No | 0.5548 |
| SlBZR1b-D b22:BRZ- vs. SlBZR1b-D b6:BRZ+ | 3.815 | 2.696 to 4.933 | Yes | <0.0001 |
| SlBZR1b-D b22:BRZ+ vs. SlBZR1b-D b6:BRZ- | -3.588 | -4.784 to -2.391 | Yes | <0.0001 |
| SlBZR1b-D b22:BRZ+ vs. SlBZR1b-D b6:BRZ+ | 0.8923 | -0.2605 to 2.045 | No | 0.2223 |
| SlBZR1b-D b6:BRZ- vs. SlBZR1b-D b6:BRZ+ | 4.480 | 3.284 to 5.676 | Yes | <0.0001 |

**Supp. Fig. 9: BRZ treatment of second *Slbzr1a-D* independent line and *Slbzr1b-D* lines. A)** Hypocotyl length of M82 WT and *Slbzr1a-D* line a10 upon BRZ treatment grown in dark for 7 days. **B)** Statistical analysis results of Panel A. **C)** Hypocotyl length of M82 WT and *Slbzr1b-D* lines b22 and b6 upon BRZ treatment grown in dark for 7 days. **D)** Statistical analysis results of Panel C.

**A**

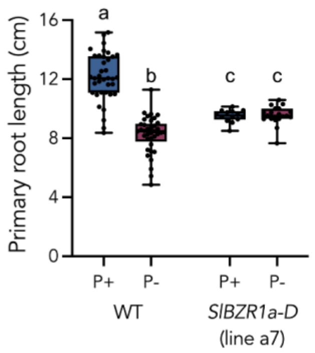

**B**

| Source of Variation | % of total variation | P value | P value summary | Significant? |
|---|---|---|---|---|
| Interaction | 19.20 | <0.0001 | **** | Yes |
| Genotype | 2.263 | 0.0124 | * | Yes |
| P treatment | 18.29 | <0.0001 | **** | Yes |
| ANOVA table | SS (Type III) | DF | MS | F (DFn, DFd) |
| Interaction | 87.28 | 1 | 87.28 | F (1, 103) = 55.00 |
| Genotype | 10.28 | 1 | 10.28 | F (1, 103) = 6.481 |
| P treatment | 83.14 | 1 | 83.14 | F (1, 103) = 52.39 |
| Residual | 163.5 | 103 | 1.587 | |

| Tukey's multiple comparisons | Predicted mean diff. | 95.00% CI of diff. | Below threshold? | Adjusted P Value |
|---|---|---|---|---|
| wt:P+ vs. wt:P- | 3.895 | 3.131 to 4.660 | Yes | <0.0001 |
| wt:P+ vs. SlBZR1a-D a7:P+ | 2.648 | 1.616 to 3.680 | Yes | <0.0001 |
| wt:P+ vs. SlBZR1a-D a7:P- | 2.601 | 1.672 to 3.529 | Yes | <0.0001 |
| wt:P- vs. SlBZR1a-D a7:P+ | -1.247 | -2.280 to -0.2151 | Yes | 0.0111 |
| wt:P- vs. SlBZR1a-D a7:P- | -1.295 | -2.223 to -0.3662 | Yes | 0.0024 |
| SlBZR1a-D a7:P+ vs. SlBZR1a-D a7:P- | -0.04729 | -1.206 to 1.111 | No | 0.9996 |

**C**

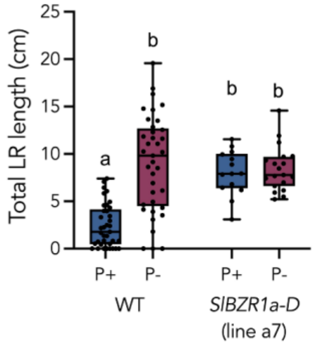

**D**

| Source of Variation | % of total variation | P value | P value summary | Significant? |
|---|---|---|---|---|
| Interaction | 9.586 | <0.0001 | **** | Yes |
| Genotype | 6.229 | 0.0012 | ** | Yes |
| P treatment | 12.25 | <0.0001 | **** | Yes |
| ANOVA table | SS (Type III) | DF | MS | F (DFn, DFd) |
| Interaction | 215.6 | 1 | 215.6 | F (1, 103) = 17.14 |
| Genotype | 140.1 | 1 | 140.1 | F (1, 103) = 11.14 |
| P treatment | 275.5 | 1 | 275.5 | F (1, 103) = 21.90 |
| Residual | 1296 | 103 | 12.58 | |

| Tukey's multiple comparisons | Predicted mean diff. | 95.00% CI of diff. | Below threshold? | Adjusted P Value |
|---|---|---|---|---|
| wt:P+ vs. wt:P- | -6.601 | -8.754 to -4.447 | Yes | <0.0001 |
| wt:P+ vs. SlBZR1a-D a7:P+ | -5.596 | -8.502 to -2.690 | Yes | <0.0001 |
| wt:P+ vs. SlBZR1a-D a7:P- | -6.000 | -8.614 to -3.386 | Yes | <0.0001 |
| wt:P- vs. SlBZR1a-D a7:P+ | 1.005 | -1.901 to 3.911 | No | 0.8032 |
| wt:P- vs. SlBZR1a-D a7:P- | 0.6007 | -2.013 to 3.215 | No | 0.9318 |
| SlBZR1a-D a7:P+ vs. SlBZR1a-D a7:P- | -0.4042 | -3.666 to 2.858 | No | 0.9882 |

**Supp. Fig. 10: RSA response of second *Slbzr1a-D* independent line (a7) to P-limiting conditions. A)** Primary root length of M82 WT and *Slbzr1a-D* line a7 under P sufficiency or deficiency. **B)** Statistical analysis results of Panel A. **C)** Total lateral root length of M82 WT and *Slbzr1a-D* line a7 under P sufficiency or deficiency. **D)** Statistical analysis results of Panel C.

**A**

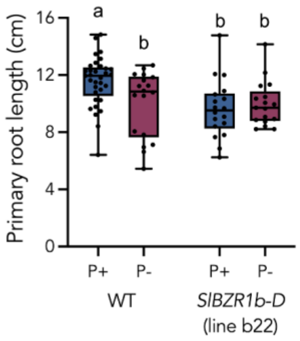

**B**

| Source of Variation | % of total variation | P value | P value summary | Significant? |
|---|---|---|---|---|
| Interaction | 5.066 | 0.0282 | * | Yes |
| Genotype | 5.404 | 0.0235 | * | Yes |
| P treatment | 2.050 | 0.1590 | ns | No |

| ANOVA table | SS (Type III) | DF | MS | F (DFn, DFd) |
|---|---|---|---|---|
| Interaction | 18.14 | 1 | 18.14 | F (1, 82) = 4.993 |
| Genotype | 19.35 | 1 | 19.35 | F (1, 82) = 5.327 |
| P treatment | 7.340 | 1 | 7.340 | F (1, 82) = 2.020 |
| Residual | 297.9 | 82 | 3.633 | |

| Tukey's multiple comparisons | Predicted mean diff. | 95.00% CI of diff. | Below threshold? | Adjusted P Value |
|---|---|---|---|---|
| WT:P+ vs. WT:P- | 1.559 | 0.09445 to 3.024 | Yes | 0.0324 |
| WT:P+ vs. SlBZR1b-D b22:P+ | 1.937 | 0.4725 to 3.402 | Yes | 0.0046 |
| WT:P+ vs. SlBZR1b-D b22:P- | 1.590 | 0.09814 to 3.083 | Yes | 0.0321 |
| WT:P- vs. SlBZR1b-D b22:P+ | 0.3781 | -1.288 to 2.044 | No | 0.9333 |
| WT:P- vs. SlBZR1b-D b22:P- | 0.03131 | -1.659 to 1.722 | No | >0.9999 |
| SlBZR1b-D b22:P+ vs. SlBZR1b-D b22:P- | -0.3467 | -2.037 to 1.344 | No | 0.9495 |

**C**

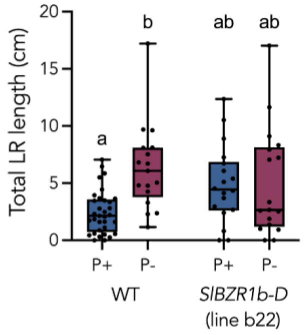

**D**

| Source of Variation | % of total variation | P value | P value summary | Significant? |
|---|---|---|---|---|
| Interaction | 6.486 | 0.0133 | * | Yes |
| Genotype | 0.5110 | 0.4792 | ns | No |
| P treatment | 7.154 | 0.0094 | ** | Yes |

| ANOVA table | SS (Type III) | DF | MS | F (DFn, DFd) |
|---|---|---|---|---|
| Interaction | 71.11 | 1 | 71.11 | F (1, 81) = 6.413 |
| Genotype | 5.602 | 1 | 5.602 | F (1, 81) = 0.5053 |
| P treatment | 78.43 | 1 | 78.43 | F (1, 81) = 7.074 |
| Residual | 898.1 | 81 | 11.09 | |

| Tukey's multiple comparisons | Predicted mean diff. | 95.00% CI of diff. | Below threshold? | Adjusted P Value |
|---|---|---|---|---|
| WT:P+ vs. WT:P- | -3.900 | -6.507 to -1.292 | Yes | 0.0010 |
| WT:P+ vs. SlBZR1b-D b22:P+ | -2.436 | -4.995 to 0.1235 | No | 0.0680 |
| WT:P+ vs. SlBZR1b-D b22:P- | -2.531 | -5.139 to 0.07615 | No | 0.0603 |
| WT:P- vs. SlBZR1b-D b22:P+ | 1.464 | -1.490 to 4.418 | No | 0.5658 |
| WT:P- vs. SlBZR1b-D b22:P- | 1.368 | -1.628 to 4.364 | No | 0.6299 |
| SlBZR1b-D b22:P+ vs. SlBZR1b-D b22:P- | -0.09557 | -3.050 to 2.858 | No | 0.9998 |

**Supp. Fig. 11: RSA response of *Slbzr1b-D* line (b22) to P-limiting conditions. A)** Primary root length of M82 WT and *Slbzr1b-D* line b22 under P sufficiency or deficiency. **B)** Statistical analysis results of Panel A. **C)** Total lateral root length of M82 WT and *Slbzr1b-D* line b22 under P sufficiency or deficiency. **D)** Statistical analysis results of Panel C.

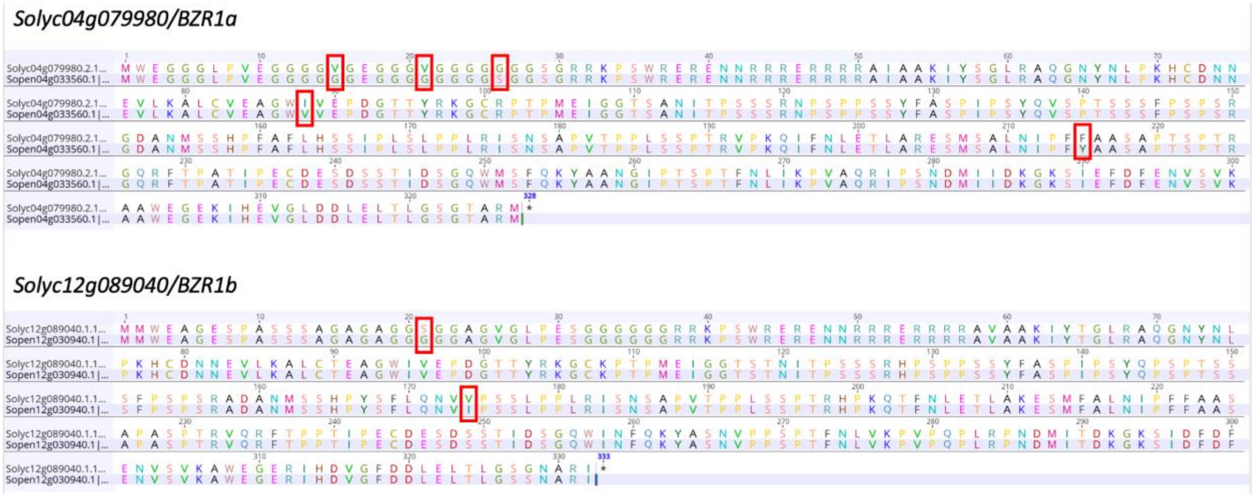

**Supp. Fig. 12: Amino acid sequence comparison of *Solyc12g089040* and *Solyc04g079980* with *S. pennellii* BZR1 orthologs shows no mutation in the predicted phosphorylation sites.**

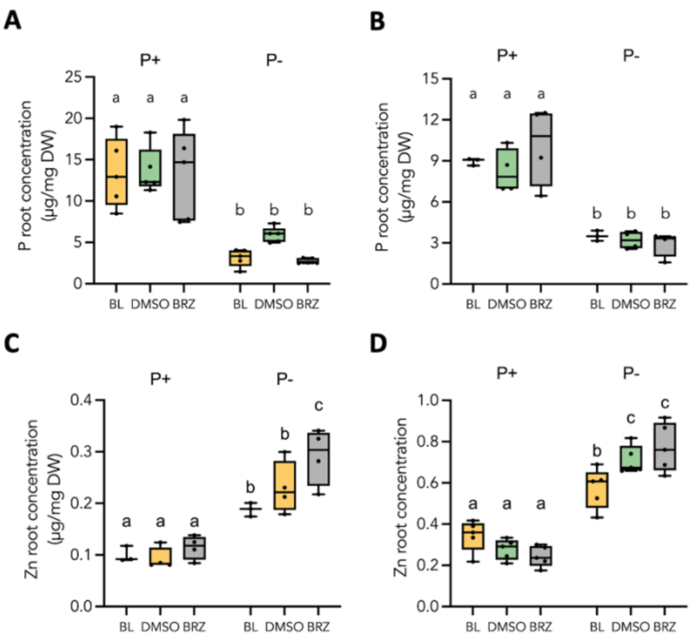

**E**

| Ion | P treatment p-value | BR treatment p-value | Interaction p-value |
|---|---|---|---|
| P | <2e-16 | 0.3349 | 0.5474 |
| Zn | <2e-16 | 0.0235 | **0.0006** |

**F**

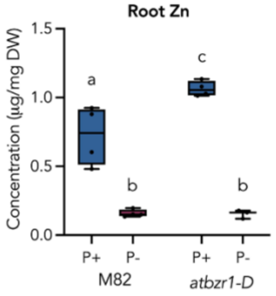

**Supp. Fig 13: Two additional replicates for M82 BL/BRZ treatment ICP-MS results. A)** Second **B)** and third replicate for P root concentration. **C)** Second **D)** and third replicate for Zn root concentration. **E)** Three independent experiments (batches) were analyzed. Thus, to determine the effect of P treatment and BRZ or BL (BR) treatment on P and Zn concentrations in the root, mixed model analysis of variance (ANOVA) was performed with P, BR and their interaction as fixed factors and the batch as random factor, followed by Correction for multiple comparisons. Post-hoc analysis was performed using pairwise differences of LS-mean as implemented in the lmerTest R package. P-values are based on the t-distribution using degrees of freedom according to Satterthwaites method. P-values that passed correction for multiple comparisons using the False Discovery Rate procedure with alpha<0.05 are indicated by letters. F) Arabidopsis *atbzr1-D* accumulates Zn in P sufficient conditions compared to Col-0 wild-type.

**A**

| Source of Variation | % of total variation | P value | P value summary | Significant? |
|---|---|---|---|---|
| Interaction | 0.05551 | 0.8495 | ns | No |
| Genotype | 10.52 | 0.0204 | * | Yes |
| P treatment | 71.69 | <0.0001 | **** | Yes |

| ANOVA table | SS (Type III) | DF | MS | F (DFn, DFd) | P value |
|---|---|---|---|---|---|
| Interaction | 0.01843 | 1 | 0.01843 | F (1, 12) = 0.03757 | P=0.8495 |
| Genotype | 3.494 | 1 | 3.494 | F (1, 12) = 7.124 | P=0.0204 |
| P treatment | 23.80 | 1 | 23.80 | F (1, 12) = 48.53 | P<0.0001 |
| Residual | 5.886 | 12 | 0.4905 | | |

| Tukey's multiple comparisons | Predicted mean diff. | 95.00% CI of diff. | Below threshold? | Adjusted P Value |
|---|---|---|---|---|
| M82:P+ vs. M82:P- | 2.507 | 0.9517 to 4.063 | Yes | 0.0017 |
| M82:P+ vs. *bzr1*:P+ | 1.003 | -0.5531 to 2.558 | No | 0.3352 |
| M82:P+ vs. *bzr1*:P- | 3.374 | 1.818 to 4.930 | Yes | 0.0001 |
| M82:P- vs. *bzr1*:P+ | -1.505 | -3.060 to 0.05085 | No | 0.0602 |
| M82:P- vs. *bzr1*:P- | 0.8668 | -0.6888 to 2.422 | No | 0.4880 |
| *bzr1*:P+ vs. *bzr1*:P- | 2.372 | 0.8159 to 3.927 | Yes | 0.0026 |

**B**

| Source of Variation | % of total variation | P value | P value summary | Significant? |
|---|---|---|---|---|
| Interaction | 0.0001080 | 0.9803 | ns | No |
| Genotype | 0.8588 | 0.0444 | * | Yes |
| P treatment | 97.10 | <0.0001 | **** | Yes |

| ANOVA table | SS (Type III) | DF | MS | F (DFn, DFd) | P value |
|---|---|---|---|---|---|
| Interaction | 3.906e-005 | 1 | 3.906e-005 | F (1, 12) = 0.0006342 | P=0.9803 |
| Genotype | 0.3105 | 1 | 0.3105 | F (1, 12) = 5.041 | P=0.0444 |
| P treatment | 35.11 | 1 | 35.11 | F (1, 12) = 570.0 | P<0.0001 |
| Residual | 0.7392 | 12 | 0.06160 | | |

| Tukey's multiple comparisons | Predicted mean diff. | 95.00% CI of diff. | Below threshold? | Adjusted P Value |
|---|---|---|---|---|
| M82:P+ vs. M82:P- | 2.960 | 2.408 to 3.511 | Yes | <0.0001 |
| M82:P+ vs. *bzr1*:P+ | 0.2755 | -0.2758 to 0.8268 | No | 0.6022 |
| M82:P+ vs. *bzr1*:P- | 3.241 | 2.690 to 3.793 | Yes | <0.0001 |
| M82:P- vs. *bzr1*:P+ | -2.684 | -3.235 to -2.133 | Yes | <0.0001 |
| M82:P- vs. *bzr1*:P- | 0.2818 | -0.2695 to 0.8330 | No | 0.5793 |
| *bzr1*:P+ vs. *bzr1*:P- | 2.966 | 2.414 to 3.517 | Yes | <0.0001 |

**C**

| Source of Variation | % of total variation | P value | P value summary | Significant? |
|---|---|---|---|---|
| Interaction | 4.459 | 0.1540 | ns | No |
| Genotype | 6.667 | 0.0875 | ns | No |
| P treatment | 65.76 | <0.0001 | **** | Yes |

| ANOVA table | SS (Type III) | DF | MS | F (DFn, DFd) | P value |
|---|---|---|---|---|---|
| Interaction | 0.009752 | 1 | 0.009752 | F (1, 12) = 2.315 | P=0.1540 |
| Genotype | 0.01458 | 1 | 0.01458 | F (1, 12) = 3.461 | P=0.0875 |
| P treatment | 0.1438 | 1 | 0.1438 | F (1, 12) = 34.14 | P<0.0001 |
| Residual | 0.05055 | 12 | 0.004212 | | |

| Tukey's multiple comparisons | Predicted mean diff. | 95.00% CI of diff. | Below threshold? | Adjusted P Value |
|---|---|---|---|---|
| M82:P+ vs. M82:P- | -0.2390 | -0.3832 to -0.09484 | Yes | 0.0013 |
| M82:P+ vs. *bzr1*:P+ | -0.1098 | -0.2539 to 0.03441 | No | 0.1877 |
| M82:P+ vs. *bzr1*:P- | -0.2500 | -0.3942 to -0.1058 | Yes | 0.0009 |
| M82:P- vs. *bzr1*:P+ | 0.1293 | -0.01491 to 0.2734 | No | 0.0898 |
| M82:P- vs. *bzr1*:P- | -0.01100 | -0.1552 to 0.1332 | No | >0.9999 |
| *bzr1*:P+ vs. *bzr1*:P- | -0.1403 | -0.2844 to 0.003913 | No | 0.0584 |

**D**

| Source of Variation | % of total variation | P value | P value summary | Significant? |
|---|---|---|---|---|
| Interaction | 8.414 | <0.0001 | **** | Yes |
| Genotype | 1.628 | 0.0091 | ** | Yes |
| P treatment | 87.93 | <0.0001 | **** | Yes |

| ANOVA table | SS (Type III) | DF | MS | F (DFn, DFd) | P value |
|---|---|---|---|---|---|
| Interaction | 3.136 | 1 | 3.136 | F (1, 12) = 49.83 | P<0.0001 |
| Genotype | 0.6068 | 1 | 0.6068 | F (1, 12) = 9.640 | P=0.0091 |
| P treatment | 32.78 | 1 | 32.78 | F (1, 12) = 520.7 | P<0.0001 |
| Residual | 0.7554 | 12 | 0.06295 | | |

| Tukey's multiple comparisons | Predicted mean diff. | 95.00% CI of diff. | Below threshold? | Adjusted P Value |
|---|---|---|---|---|
| M82:P+ vs. M82:P- | -3.748 | -4.305 to -3.191 | Yes | <0.0001 |
| M82:P+ vs. *bzr1*:P+ | -0.4960 | -1.053 to 0.06128 | No | 0.0932 |
| M82:P+ vs. *bzr1*:P- | -2.473 | -3.030 to -1.916 | Yes | <0.0001 |
| M82:P- vs. *bzr1*:P+ | 3.252 | 2.695 to 3.809 | Yes | <0.0001 |
| M82:P- vs. *bzr1*:P- | 1.275 | 0.7177 to 1.832 | Yes | <0.0001 |
| *bzr1*:P+ vs. *bzr1*:P- | -1.977 | -2.534 to -1.420 | Yes | <0.0001 |

**E**

| Source of Variation | % of total variation | P value | P value summary | Significant? |
|---|---|---|---|---|
| Interaction | 65.00 | <0.0001 | **** | Yes |
| Genotype | 13.76 | 0.0161 | * | Yes |
| P treatment | 0.1604 | 0.7677 | ns | No |

| ANOVA table | SS (Type III) | DF | MS | F (DFn, DFd) | P value |
|---|---|---|---|---|---|
| Interaction | 0.1206 | 1 | 0.1206 | F (1, 12) = 37.01 | P<0.0001 |
| Genotype | 0.02552 | 1 | 0.02552 | F (1, 12) = 7.832 | P=0.0161 |
| P treatment | 0.0002976 | 1 | 0.0002976 | F (1, 12) = 0.09132 | P=0.7677 |
| Residual | 0.03910 | 12 | 0.003258 | | |

| Tukey's multiple comparisons | Predicted mean diff. | 95.00% CI of diff. | Below threshold? | Adjusted P Value |
|---|---|---|---|---|
| M82:P+ vs. M82:P- | -0.1650 | -0.2918 to -0.03821 | Yes | 0.0090 |
| M82:P+ vs. *bzr1*:P+ | -0.2535 | -0.3803 to -0.1267 | Yes | 0.0002 |
| M82:P+ vs. *bzr1*:P- | -0.07125 | -0.1980 to 0.05554 | No | 0.4789 |
| M82:P- vs. *bzr1*:P+ | -0.08850 | -0.2153 to 0.03829 | No | 0.2592 |
| M82:P- vs. *bzr1*:P- | 0.09375 | -0.03304 to 0.2205 | No | 0.2103 |
| *bzr1*:P+ vs. *bzr1*:P- | 0.1823 | 0.05546 to 0.3090 | Yes | 0.0042 |

**F**

| Source of Variation | % of total variation | P value | P value summary | Significant? |
|---|---|---|---|---|
| Interaction | 81.53 | <0.0001 | **** | Yes |
| Genotype | 1.686 | 0.1288 | ns | No |
| P treatment | 9.174 | 0.0025 | ** | Yes |

| ANOVA table | SS (Type III) | DF | MS | F (DFn, DFd) | P value |
|---|---|---|---|---|---|
| Interaction | 22.50 | 1 | 22.50 | F (1, 12) = 128.7 | P<0.0001 |
| Genotype | 0.4651 | 1 | 0.4651 | F (1, 12) = 2.660 | P=0.1288 |
| P treatment | 2.531 | 1 | 2.531 | F (1, 12) = 14.48 | P=0.0025 |
| Residual | 2.098 | 12 | 0.1749 | | |

| Tukey's multiple comparisons | Predicted mean diff. | 95.00% CI of diff. | Below threshold? | Adjusted P Value |
|---|---|---|---|---|
| M82:P+ vs. M82:P- | -1.576 | -2.505 to -0.6472 | Yes | 0.0011 |
| M82:P+ vs. *bzr1*:P+ | -2.031 | -2.959 to -1.102 | Yes | 0.0001 |
| M82:P+ vs. *bzr1*:P- | 1.137 | 0.2077 to 2.065 | Yes | 0.0139 |
| M82:P- vs. *bzr1*:P+ | -0.4545 | -1.383 to 0.4743 | No | 0.6234 |
| M82:P- vs. *bzr1*:P- | 2.713 | 1.784 to 3.641 | Yes | <0.0001 |
| *bzr1*:P+ vs. *bzr1*:P- | 3.167 | 2.238 to 4.096 | Yes | <0.0001 |

**Supp. Fig 14: Statistical analysis of Figure 7. A)** Figure 7 Panel C (Root P). **B)** Figure 7 Panel D (Shoot P). **C)** Figure 7 Panel E (Root Fe). **D)** Figure 7 Panel F (Shoot Fe). **E)** Figure 7 Panel G (Root Zn). **F)** Figure 7 Panel H (Shoot Zn).

**A**

| Source of Variation | % of total variation | P value | P value summary | Significant? |
|---|---|---|---|---|
| Interaction | 1.768 | 0.1072 | ns | No |
| Zn treatment | 9.510 | 0.0003 | *** | Yes |
| P treatment | 24.53 | <0.0001 | **** | Yes |

| ANOVA table | SS (Type III) | DF | MS | F (DFn, DFd) | P value |
|---|---|---|---|---|---|
| Interaction | 40.08 | 1 | 40.08 | F (1, 96) = 2.644 | P=0.1072 |
| Zn treatment | 215.6 | 1 | 215.6 | F (1, 96) = 14.22 | P=0.0003 |
| P treatment | 556.2 | 1 | 556.2 | F (1, 96) = 36.69 | P<0.0001 |
| Residual | 1455 | 96 | 15.16 | | |

| Tukey's multiple comparisons | Predicted mean diff. | 95.00% CI of diff. | Below threshold? | Adjusted P Value |
|---|---|---|---|---|
| Zn+:P+ vs. Zn+:P- | 5.983 | 3.103 to 8.862 | Yes | <0.0001 |
| Zn+:P+ vs. Zn-:P+ | 4.203 | 1.324 to 7.082 | Yes | 0.0014 |
| Zn+:P+ vs. Zn-:P- | 7.653 | 4.774 to 10.53 | Yes | <0.0001 |
| Zn+:P- vs. Zn-:P+ | -1.780 | -4.659 to 1.100 | No | 0.3745 |
| Zn+:P- vs. Zn-:P- | 1.671 | -1.209 to 4.550 | No | 0.4314 |
| Zn-:P+ vs. Zn-:P- | 3.450 | 0.5711 to 6.330 | Yes | 0.0121 |

**B**

| Source of Variation | % of total variation | P value | P value summary | Significant? |
|---|---|---|---|---|
| Interaction | 0.3818 | 0.4424 | ns | No |
| Zn treatment | 2.446 | 0.0536 | ns | No |
| P treatment | 27.54 | <0.0001 | **** | Yes |

| ANOVA table | SS (Type III) | DF | MS | F (DFn, DFd) | P value |
|---|---|---|---|---|---|
| Interaction | 7.477 | 1 | 7.477 | F (1, 108) = 0.5945 | P=0.4424 |
| Zn treatment | 47.91 | 1 | 47.91 | F (1, 108) = 3.809 | P=0.0536 |
| P treatment | 539.3 | 1 | 539.3 | F (1, 108) = 42.88 | P<0.0001 |
| Residual | 1358 | 108 | 12.58 | | |

| Tukey's multiple comparisons | Predicted mean diff. | 95.00% CI of diff. | Below threshold? | Adjusted P Value |
|---|---|---|---|---|
| Zn+:P+ vs. Zn+:P- | -3.874 | -6.393 to -1.356 | Yes | 0.0006 |
| Zn+:P+ vs. Zn-:P+ | -0.7918 | -3.267 to 1.683 | No | 0.8378 |
| Zn+:P+ vs. Zn-:P- | -5.700 | -8.175 to -3.225 | Yes | <0.0001 |
| Zn+:P- vs. Zn-:P+ | 3.083 | 0.6077 to 5.558 | Yes | 0.0083 |
| Zn+:P- vs. Zn-:P- | -1.826 | -4.301 to 0.6490 | No | 0.2236 |
| Zn-:P+ vs. Zn-:P- | -4.909 | -7.339 to -2.478 | Yes | <0.0001 |

**C**

| Source of Variation | % of total variation | P value | P value summary | Significant? |
|---|---|---|---|---|
| Interaction | 0.8841 | 0.0085 | ** | Yes |
| Zn treatment | 1.271 | 0.0016 | ** | Yes |
| P treatment | 47.76 | <0.0001 | **** | Yes |

| ANOVA table | SS (Type III) | DF | MS | F (DFn, DFd) | P value |
|---|---|---|---|---|---|
| Interaction | 0.8273 | 1 | 0.8273 | F (1, 396) = 6.991 | P=0.0085 |
| Zn treatment | 1.190 | 1 | 1.190 | F (1, 396) = 10.05 | P=0.0016 |
| P treatment | 44.69 | 1 | 44.69 | F (1, 396) = 377.7 | P<0.0001 |
| Residual | 46.87 | 396 | 0.1183 | | |

| Tukey's multiple comparisons | Predicted mean diff. | 95.00% CI of diff. | Below threshold? | Adjusted P Value |
|---|---|---|---|---|
| Zn+:P+ vs. Zn+:P- | -0.7595 | -0.8850 to -0.6340 | Yes | <0.0001 |
| Zn+:P+ vs. M82:P+ | 0.01812 | -0.1074 to 0.1436 | No | 0.9824 |
| Zn+:P+ vs. M82:P- | -0.5595 | -0.6850 to -0.4339 | Yes | <0.0001 |
| Zn+:P- vs. M82:P+ | 0.7776 | 0.6521 to 0.9031 | Yes | <0.0001 |
| Zn+:P- vs. M82:P- | 0.2000 | 0.07451 to 0.3256 | Yes | 0.0003 |
| M82:P+ vs. M82:P- | -0.5776 | -0.7031 to -0.4521 | Yes | <0.0001 |

**D**

| Source of Variation | % of total variation | P value | P value summary | Significant? |
|---|---|---|---|---|
| Interaction | 0.8823 | 0.0108 | * | Yes |
| Zn treatment | 0.4410 | 0.0690 | ns | No |
| P treatment | 86.16 | <0.0001 | **** | Yes |

| ANOVA table | SS (Type III) | DF | MS | F (DFn, DFd) | P value |
|---|---|---|---|---|---|
| Interaction | 342260 | 1 | 342260 | F (1, 96) = 6.765 | P=0.0108 |
| Zn treatment | 171090 | 1 | 171090 | F (1, 96) = 3.382 | P=0.0690 |
| P treatment | 33422620 | 1 | 33422620 | F (1, 96) = 660.6 | P<0.0001 |
| Residual | 4856884 | 96 | 50593 | | |

| Tukey's multiple comparisons | Predicted mean diff. | 95.00% CI of diff. | Below threshold? | Adjusted P Value |
|---|---|---|---|---|
| Zn+:P+ vs. Zn+:P- | -1273 | -1440 to -1107 | Yes | <0.0001 |
| Zn+:P+ vs. M82:P+ | -34.28 | -200.6 to 132.1 | No | 0.9493 |
| Zn+:P+ vs. M82:P- | -1074 | -1240 to -907.2 | Yes | <0.0001 |
| Zn+:P- vs. M82:P+ | 1239 | 1073 to 1405 | Yes | <0.0001 |
| Zn+:P- vs. M82:P- | 199.7 | 33.39 to 366.1 | Yes | 0.0119 |
| M82:P+ vs. M82:P- | -1039 | -1206 to -872.9 | Yes | <0.0001 |

**E**

| Source of Variation | % of total variation | P value | P value summary | Significant? |
|---|---|---|---|---|
| Interaction | 7.709 | 0.0011 | ** | Yes |
| Zn treatment | 18.53 | <0.0001 | **** | Yes |
| P treatment | 8.611 | 0.0006 | *** | Yes |

| ANOVA table | SS (Type III) | DF | MS | F (DFn, DFd) | P value |
|---|---|---|---|---|---|
| Interaction | 137.7 | 1 | 137.7 | F (1, 96) = 11.36 | P=0.0011 |
| Zn treatment | 330.9 | 1 | 330.9 | F (1, 96) = 27.30 | P<0.0001 |
| P treatment | 153.8 | 1 | 153.8 | F (1, 96) = 12.69 | P=0.0006 |
| Residual | 1164 | 96 | 12.12 | | |

| Tukey's multiple comparisons | Predicted mean diff. | 95.00% CI of diff. | Below threshold? | Adjusted P Value |
|---|---|---|---|---|
| Zn+:P+ vs. Zn+:P- | 0.1335 | -2.441 to 2.708 | No | 0.9991 |
| Zn+:P+ vs. Zn-:P+ | 1.291 | -1.283 to 3.866 | No | 0.5579 |
| Zn+:P+ vs. Zn-:P- | 6.118 | 3.544 to 8.693 | Yes | <0.0001 |
| Zn+:P- vs. Zn-:P+ | 1.158 | -1.417 to 3.732 | No | 0.6434 |
| Zn+:P- vs. Zn-:P- | 5.985 | 3.410 to 8.559 | Yes | <0.0001 |
| Zn-:P+ vs. Zn-:P- | 4.827 | 2.252 to 7.401 | Yes | <0.0001 |

**F**

| Source of Variation | % of total variation | P value | P value summary | Significant? |
|---|---|---|---|---|
| Interaction | 7.535 | 0.0021 | ** | Yes |
| Zn treatment | 3.619 | 0.0314 | * | Yes |
| P treatment | 6.568 | 0.0040 | ** | Yes |

| ANOVA table | SS (Type III) | DF | MS | F (DFn, DFd) | P value |
|---|---|---|---|---|---|
| Interaction | 89.48 | 1 | 89.48 | F (1, 108) = 9.901 | P=0.0021 |
| Zn treatment | 42.98 | 1 | 42.98 | F (1, 108) = 4.755 | P=0.0314 |
| P treatment | 78.00 | 1 | 78.00 | F (1, 108) = 8.630 | P=0.0040 |
| Residual | 976.1 | 108 | 9.038 | | |

| Tukey's multiple comparisons | Predicted mean diff. | 95.00% CI of diff. | Below threshold? | Adjusted P Value |
|---|---|---|---|---|
| Zn+:P+ vs. Zn+:P- | 0.1187 | -1.997 to 2.235 | No | 0.9989 |
| Zn+:P+ vs. Zn-:P+ | 0.5489 | -1.549 to 2.647 | No | 0.9035 |
| Zn+:P+ vs. Zn-:P- | -2.909 | -5.025 to -0.7929 | Yes | 0.0028 |
| Zn+:P- vs. Zn-:P+ | 0.4302 | -1.648 to 2.509 | No | 0.9490 |
| Zn+:P- vs. Zn-:P- | -3.028 | -5.124 to -0.9309 | Yes | 0.0015 |
| Zn-:P+ vs. Zn-:P- | -3.458 | -5.536 to -1.379 | Yes | 0.0002 |

**G**

| Source of Variation | % of total variation | P value | P value summary | Significant? |
|---|---|---|---|---|
| Interaction | 10.17 | <0.0001 | **** | Yes |
| Zn treatment | 24.23 | <0.0001 | **** | Yes |
| P treatment | 14.81 | <0.0001 | **** | Yes |

| ANOVA table | SS (Type III) | DF | MS | F (DFn, DFd) | P value |
|---|---|---|---|---|---|
| Interaction | 5.105 | 1 | 5.105 | F (1, 396) = 79.35 | P<0.0001 |
| Zn treatment | 12.16 | 1 | 12.16 | F (1, 396) = 189.0 | P<0.0001 |
| P treatment | 7.433 | 1 | 7.433 | F (1, 396) = 115.5 | P<0.0001 |
| Residual | 25.48 | 396 | 0.06433 | | |

| Tukey's multiple comparisons | Predicted mean diff. | 95.00% CI of diff. | Below threshold? | Adjusted P Value |
|---|---|---|---|---|
| Zn+:P+ vs. Zn+:P- | 0.4986 | 0.4060 to 0.5911 | Yes | <0.0001 |
| Zn+:P+ vs. bzr1:P+ | 0.5746 | 0.4821 to 0.6672 | Yes | <0.0001 |
| Zn+:P+ vs. bzr1:P- | 0.6213 | 0.5288 to 0.7139 | Yes | <0.0001 |
| Zn+:P- vs. bzr1:P+ | 0.07607 | -0.01648 to 0.1686 | No | 0.1483 |
| Zn+:P- vs. bzr1:P- | 0.1228 | 0.03021 to 0.2153 | Yes | 0.0038 |
| bzr1:P+ vs. bzr1:P- | 0.04669 | -0.04586 to 0.1392 | No | 0.5624 |

**H**

| Source of Variation | % of total variation | P value | P value summary | Significant? |
|---|---|---|---|---|
| Interaction | 0.4573 | 0.0070 | ** | Yes |
| Zn treatment | 0.08228 | 0.2456 | ns | No |
| P treatment | 93.67 | <0.0001 | **** | Yes |

| ANOVA table | SS (Type III) | DF | MS | F (DFn, DFd) | P value |
|---|---|---|---|---|---|
| Interaction | 787479 | 1 | 787479 | F (1, 96) = 7.584 | P=0.0070 |
| Zn treatment | 141677 | 1 | 141677 | F (1, 96) = 1.365 | P=0.2456 |
| P treatment | 161300160 | 1 | 161300160 | F (1, 96) = 1554 | P<0.0001 |
| Residual | 9967504 | 96 | 103828 | | |

| Tukey's multiple comparisons | Predicted mean diff. | 95.00% CI of diff. | Below threshold? | Adjusted P Value |
|---|---|---|---|---|
| Zn+:P+ vs. Zn+:P- | -2363 | -2601 to -2124 | Yes | <0.0001 |
| Zn+:P+ vs. Zn-:P+ | 252.8 | 14.47 to 491.1 | Yes | 0.0332 |
| Zn+:P+ vs. Zn-:P- | -2465 | -2703 to -2227 | Yes | <0.0001 |
| Zn+:P- vs. Zn-:P+ | 2615 | 2377 to 2854 | Yes | <0.0001 |
| Zn+:P- vs. Zn-:P- | -102.2 | -340.5 to 136.1 | No | 0.6773 |
| Zn-:P+ vs. Zn-:P- | -2718 | -2956 to -2479 | Yes | <0.0001 |

**Supp. Fig 15: Statistical analysis of Figure 8 of A)** Primary root length **B)** Total lateral root length **C)** Mature root hair length and **D)** APase activity of M82 WT in P-sufficient or P-deficient conditions when Zn is present or absent. **E)** Primary root length **F)** Total lateral root length **G)** Mature root hair length and **H)** APase activity of *Slbzr1a-D* in P-sufficient or P-deficient conditions when Zn is present or absent.
